# Kinematically-Dominated Regime Shapes Cell Traction Force Dynamics under Osmotic Shock

**DOI:** 10.64898/2026.09.21.715587

**Authors:** Jiarui Gan, Xiapeng Wang, Wenjie Wu, Qingchuan Zhang, Shubo Zhang, Shangquan Wu

## Abstract

Adherent cells must continuously adapt to rapid environmental fluctuations to preserve mechanical integrity. However, conventional theoretical frameworks predominantly rely on quasi-static assumptions, thereby limiting their ability to capture the transient dynamics of cellular responses to high-rate perturbations, such as acute osmotic shocks. To address this limitation, we developed a biophysical model incorporating a Hill-type law that couples active cytoskeletal mechanics to cell-edge velocity, explaining the counterintuitive experimental observation that traction forces transiently decrease during rapid hypotonic swelling despite an increase in cell size. We further construct a kinematic-geometric phase diagram that connects classical quasi-static theories with our dynamic model. Moreover, we show that traction-force dynamics are sensitive to the loading rate and amplitude of osmotic shock. In addition, stiffness-associated simulations and supporting perturbation evidence suggest a cellular stiffness-associated recovery trend in which stiffer cells exhibit faster volume recovery driven by stronger recoil of hydrostatic pressure.

**Statement of Significance:** Adherent cells often encounter sudden osmotic changes while remaining mechanically connected to their surroundings. During such rapid perturbations, traction forces may change in ways that cannot be inferred from cell size alone. This study develops a dynamic mechanical framework that links osmotic transport, cell-edge motion, cytoskeletal mechanics, and adhesion-mediated traction. The resulting regime map helps distinguish when traction is governed mainly by adhesion geometry and when it is shaped by rapid edge motion. This framework provides a testable baseline for interpreting transient traction-force responses and for extending quasi-static descriptions of adherent-cell mechanics to dynamic perturbations.

## 1 Introduction

The mechanical microenvironment of adherent cells is rarely static; instead, it is characterized by continuous, rapid fluctuations that drive cellular systems far from equilibrium (1–7). From pulsatile blood flow to traumatic injury, cells must constantly adapt to high-rate mechanical perturbations to maintain homeostasis (8, 9). Among these, acute osmotic shock serves as a paradigmatic example of a rapid volumetric perturbation (10–12), capable of instantaneously disrupting the delicate mechano-chemical equilibrium within the cytoskeleton. This disruption triggers a spectrum of responses, ranging from volume regulation and directed migration (e.g., during cancer invasion) to pathological damage under excessive loads (13, 14). Crucially, unlike suspended cells which can swell or shrink freely as isolated pressurized capsules, most cells within biological tissues must cope with these volumetric fluctuations while remaining physically tethered to the extracellular matrix (ECM). Any osmotic-induced volume change transmits deformation to the underlying ECM (15). From a fundamental mechanics perspective, such deformation is intrinsically coupled with stress re-distribution (16). Consequently, osmotic shocks do not merely alter cell size but compel a dynamic evolution of cell traction forces. These fluctuations in cell-ECM mechanics are not trivial; aberrant force transmission can induce tissue lesions and has been implicated in the pathogenesis of various diseases (17–21).

A growing body of experimental research has attempted to decouple the mechanical regulation of osmotic processes, driven by advanced micromanipulation and imaging techniques. At the cellular level, atomic force microscopy (AFM) and confocal microscopy have proven instrumental in quantifying intracellular hydrostatic pressure and 3D volume dynamics, respectively, revealing a rapid self-recovery mechanism dependent on the actomyosin cortex (22, 23). Simultaneously, at the cell-substrate interface, traction force microscopy (TFM) has demonstrated that osmotic pressure modulates traction forces (15, 24). However, despite these methodological strides, a fundamental challenge remains: distinguishing between passive mechanical deformation (e.g., osmotic swelling) and active biochemical regulation (e.g., cytoskeletal remodeling), as these events often occur concomitantly (25). Furthermore, the complex boundary conditions introduced by dynamic traction forces obscure the fundamental relationship between intracellular pressure and cortical stress, necessitating a theoretical framework to isolate these coupled interactions (26, 27).

Therefore, theoretical modeling has emerged as an indispensable tool to decouple these multifaceted interactions, offering insights into the feedback mechanisms that are difficult to isolate experimentally (28). Theoretical frameworks for suspended cells have matured from fundamental volume regulation to complex migration mechanisms. Building on Hoffman’s pump-leak model (10), Jiang integrated mechanosensitive channels with cortical mechanics to capture the coupled volumetric and mechanical responses (29). Expanding on this hydraulic perspective, Stroka and Jiang proposed the “Osmotic Engine Model”, demonstrating that directed trans-membrane water flux alone can drive cell migration in confined spaces (13). However, in vivo, most cells adhere to the extracellular matrix (ECM). This adherent state differs fundamentally from that of suspended cells due to the presence of adhesion constraints (30). Existing theoretical studies on adherent cells have primarily focused on quasi-static processes such as spreading or migration (31–36), failing to capture transient mechanical responses under dynamic loading conditions such as osmotic shock, let alone distinguish the critical roles of loading rate and magnitude. Given that the cell physically contends with such environmental fluctuations through its adhesions, traction forces emerge as the critical dynamic variable reflecting the evolution of internal stress states. However, a framework capable of predicting this spatiotemporal evolution remains elusive.

To address this gap, we developed a biophysical model that couples transmembrane transport, cytoskeletal mechanics, and adhesion-mediated traction in adherent cells under osmotic shock. The consistency between the model and osmotic recovery kinetics provides a basis for relating volume regulation to traction-force dynamics. We further use the model to examine how cell-edge velocity and adhesion geometry compete to shape traction evolution, and how the magnitude and loading rate of osmotic shock affect this process. This mechanistic interpretation also helps assess how cortical mechanical state may contribute to osmotic recovery kinetics.

## 2 Theoretical Framework

The cell is idealized as an axisymmetric, deformable body interacting with the substrate and osmotic environment. As depicted in Fig. 1, the cell is modeled as a semi-permeable continuum. Under hypotonic shock, the cell expands from its initial homeostatic state (dashed line) to a swollen state (solid line). This volume response is coupled to three transmembrane fluxes: water influx via aquaporins (*J*_*w*_), active ion pumping (*J*_*a*_) (12), and passive ion efflux (*J*_*p*_) (29). This geometric expansion induces an instantaneous remodeling of cell mechanics and energy, the formulation of which is detailed subsequently. Simultaneously, the cell exerts dynamic contractile traction forces on the substrate (red arrows), which have been shown to attenuate during hypotonic swelling (15).

**Figure 1:**
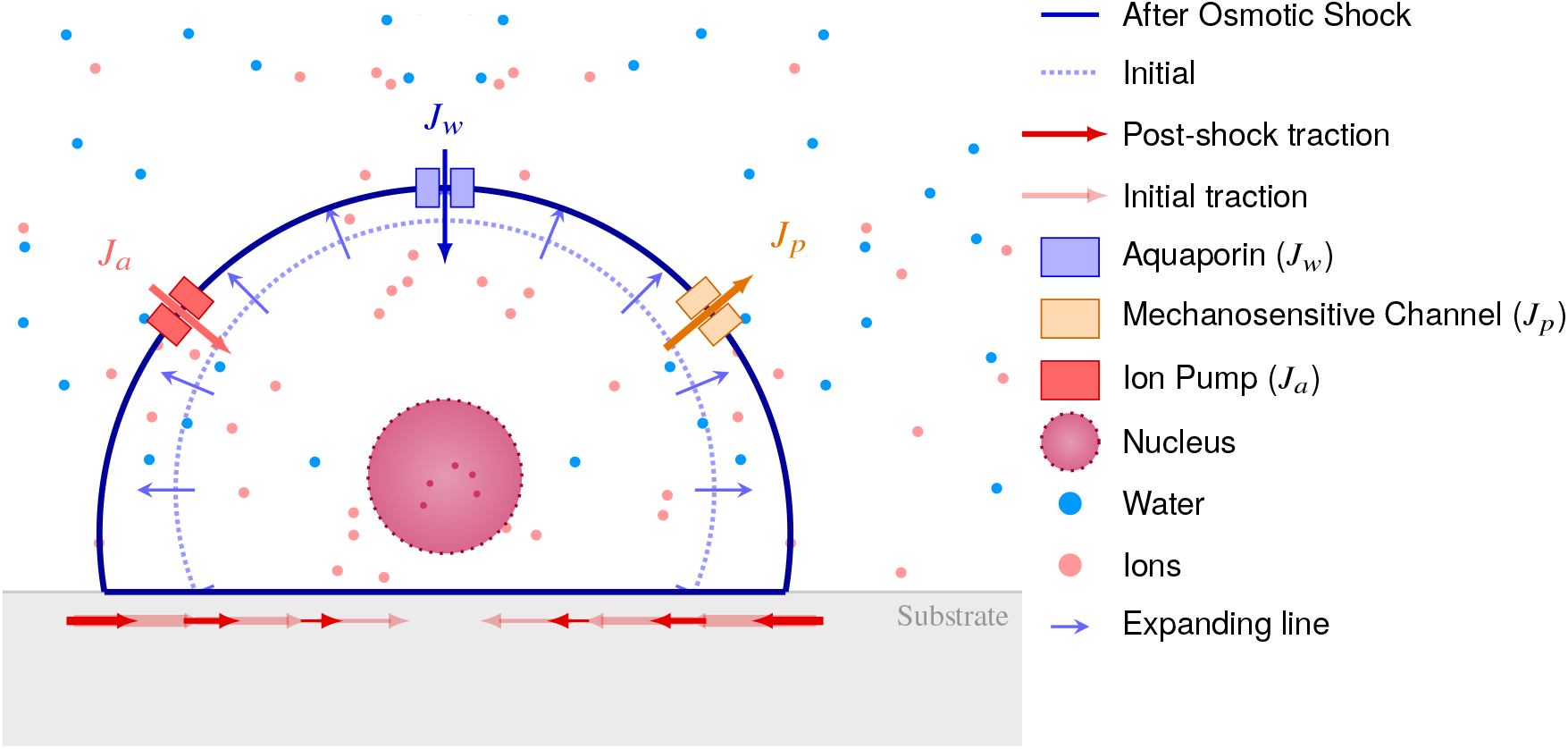
Schematic of adherent cell osmotic response dynamics. Under hypotonic shock, the cell expands from its initial homeostatic state (dashed line) to a swollen state (solid line). Volume regulation is coupled to water influx (*J*_*w*_), active ion pumping (*J*_*a*_), and passive ion efflux (*J*_*p*_), together with contractile traction forces (red arrows).

### 2.1 Geometry and Mechanics

As illustrated in Fig. 2, the adherent cell is modeled as a spherical cap of height *h* and radius *R*, adhering to the substrate over a circular contact area with adhesion radius *r*_*c*_. The geometric state can be determined by the generalized coordinates *{R, h}*(37–39). The cell volume *V*, effective surface area *A* (excluding the adhesion area), and adhesion radius *r*_*c*_ are given by:

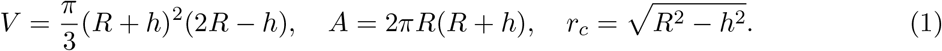

With the geometric framework established, determining the initial steady state entails the minimization of the total free energy *U* (40, 41), *whereas the subsequent mechanical evolution follows from the power balance equation. Crucially, both formulations rely on the definition of U*, which comprises four distinct components:

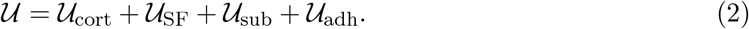

**Figure 2:**
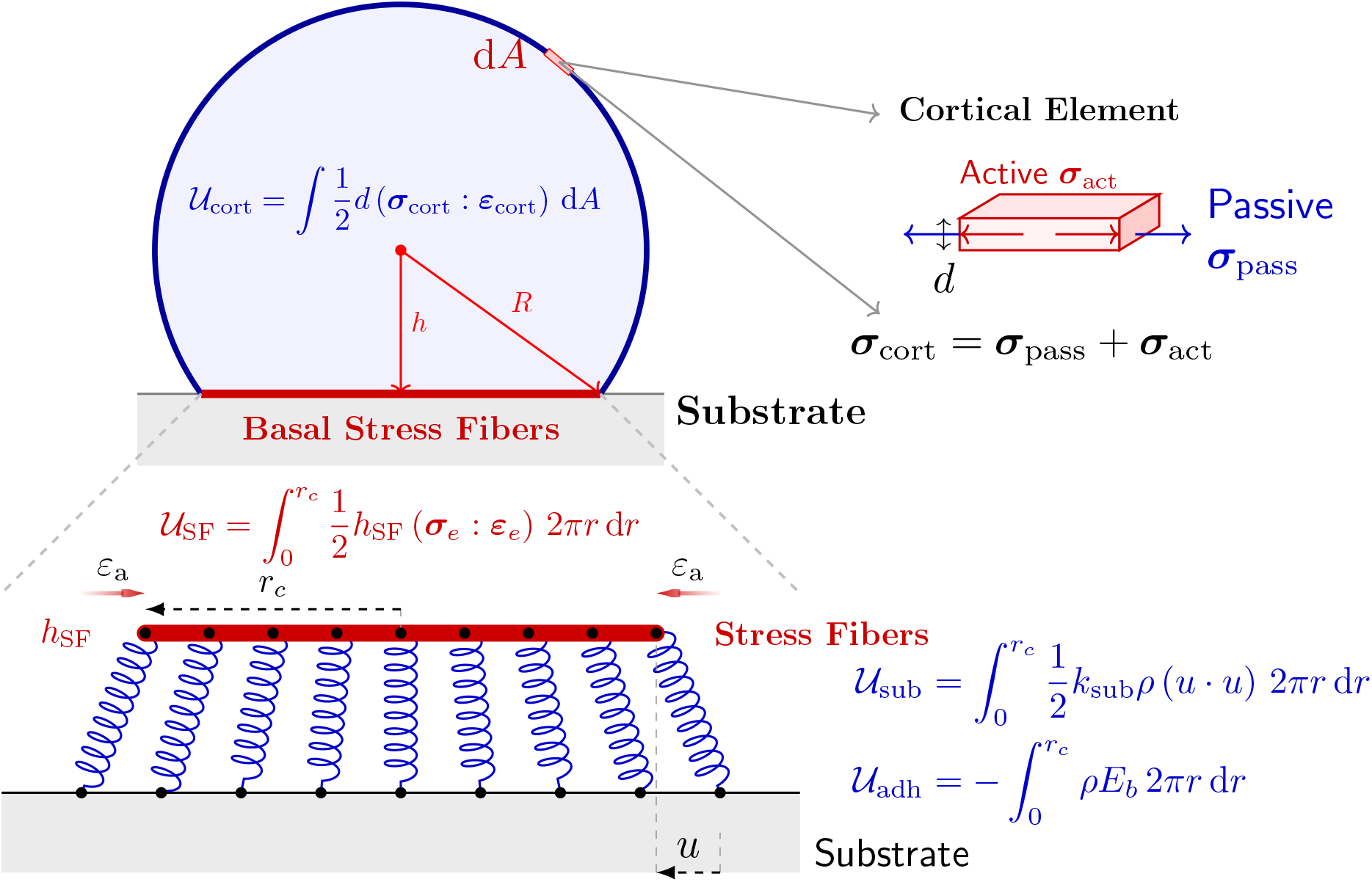
Schematic representation of the biophysical model for an adherent cell. The cell is modeled as a spherical cap characterized by the curvature radius *R* and geometric height *h*. The equilibrium state is obtained by minimizing the total free energy, *U* = *U*_SF_ + *U*_sub_ + *U*_cort_ + *U*_adh_. Specifically, the cortex (*U*_cort_) is treated as an active elastic shell generating biaxial contractile stress. The basal stress fiber network (*U*_SF_) is conceptualized as an elastic disk subject to active strain *ε*_a_ within the adhesion radius *r*_*c*_. This active contractility drives inward displacement *u*, storing elastic potential energy in the cell-substrate linkages (*U*_sub_), while the chemical energy reduction from integrin bond formation is denoted by *U*_adh_.

Here, *U*_cort_ denotes the cortical stress energy, *U*_SF_ represents the stress fiber network energy, and *U*_sub_ and *U*_adh_ correspond to the elastic and chemical energies of adhesion, respectively.

#### 2.1.1 Cortical Stress Energy (*U*_cort_)

Treating the cell cortex as an elastic thin shell, the stress energy *U*_cort_ is defined as:

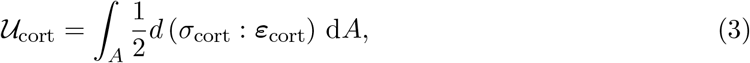

where ***ε***_cort_ is the cortical strain, and *σ*_cort_ is the surface stress determined by Young’s modulus of the cortical layer *E*_*m*_, Poisson’s ratio *ν*_*m*_, and the cortex active stress *σ*_act_:

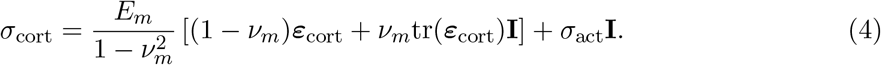

#### 2.1.2 Stress fiber network Energy (*U*_SF_)

The active contractility generated by the basal stress fiber (SF) network is modeled as an equivalent active-strained disk (42). This active contraction drives the disk inward, inducing relative displacements *u* between the integrins and the substrate, which in turn gives rise to traction forces. Defining a polar coordinate system (*r, θ*) and assuming axisymmetric deformation (negligible shear strains), the principal radial (*ε*_*r*_) and circumferential (*ε*_*θ*_) strains are:

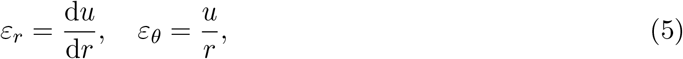

Modeling the SF network as an axisymmetric elastic continuum under plane stress, we incorporate a radial active strain *ε*_a_ to account for contractility. The resulting constitutive relations are (42):

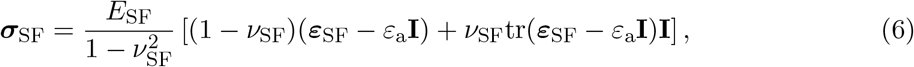

where *E*_SF_ and *ν*_SF_ denote the Young’s modulus and Poisson’s ratio of the SF network, respectively. The mechanical equilibrium of the SF network is described by:

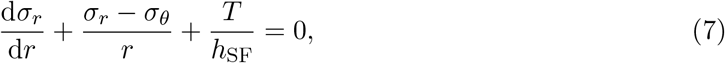

where *h*_SF_ denotes the network thickness. The traction force *T* is coupled to *u* via the adhesion density *ρ* and an effective stiffness *k*_sub_ as:

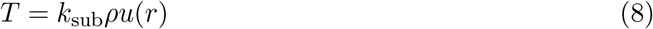

where *k*_sub_ represents the stiffness of the molecular bonds (*k*_*b*_) and the substrate (*k*_*s*_) in series (43):

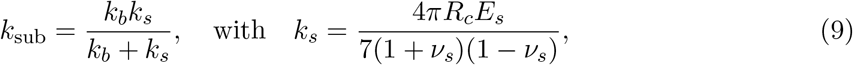

where *E*_*s*_ and *ν*_*s*_ denote the Young’s modulus and Poisson’s ratio of the substrate, respectively, and *R*_*c*_ is the effective radius of the force applied by a single bond.

Substituting the constitutive laws into the equilibrium equation Eq. (7) yields the balance equation for *u*. Assuming a radially homogeneous active strain (*∂*_*r*_*ε*_*a*_ = 0), the active stress gradients vanish:

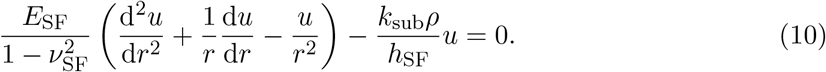

Rearranging the terms yields a modified Bessel equation:

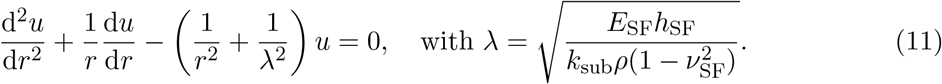

Assuming that discrete focal adhesions act collectively as an effective continuous medium over the stress transmission scale, we adopt a mean-field approximation by replacing the local density *ρ*(*r*) with its initial uniform value *ρ*_0_. This simplification allows us to interpret *λ* as a constant screening length for the shear lag effect. Equation (11) is the Bessel equation of order one, and its general solution is given by the linear combination:

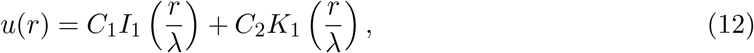

where *I*_1_ and *K*_1_ are the Bessel functions of the first and second kind, respectively. However, the function *K*_1_(*r/λ*) becomes singular (approaches infinity) as *r →* 0. To satisfy the regularity condition (bounded displacement) at the cell center, we must set *C*_2_ = 0. Consequently, the physical solution reduces to *u*(*r*) = *C*_1_*I*_1_(*r/λ*). To determine the integration constant *C*_1_, we apply the stress-free boundary condition at the cell edge (*r* = *r*_*c*_), i.e., *σ*_*r*_(*r*_*c*_) = 0. This boundary condition couples the active strain to the deformation field:

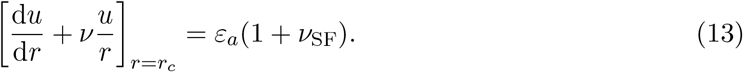

Substituting Eqs. (6) and (12) into Eq. (13) and using the derivative identity *I*_1_^*′*^ (*z*) = *I*_0_(*z*) *− I*_1_(*z*)*/z*, we obtain the final explicit expression for the displacement field:

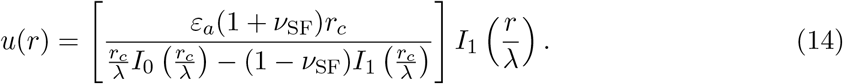

Substituting the displacement field from Eq. (14) into Eqs. (5) and (6), the stored elastic energy *U*_SF_ is obtained by integrating over the contact domain:

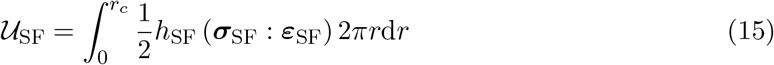

#### 2.1.3 Adhesion Elastic Energy (*U*_sub_)

During cell spreading, the contact between the cell and the substrate induces elastic energy. Modeling the integrin-substrate linkage as a series of elastic springs, we calculate the adhesion elastic energy as:

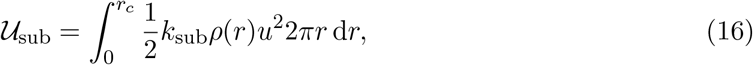

The local force acting on the bonds is determined by the relative displacement *u*(*r*). Adopting the convention that inward contraction is positive, we have:

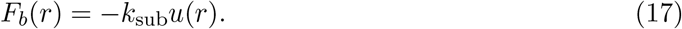

This mechanical load regulates the bond dissociation rate *k*_off_. Specifically, we employ a reduced two-pathway Bell-type catch-slip law to capture biphasic force-dependent dissociation. The parameterization follows adhesive-cell mechanics models that use catch-slip adhesion kinetics (38):

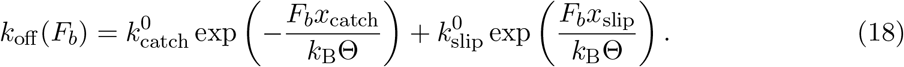

Here, *k*_B_ is the Boltzmann constant and Θ is the absolute temperature. 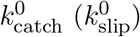 and *x*_catch_ (*x*_slip_) denote the intrinsic dissociation rates and characteristic length scales for the catch (slip) pathways, respectively. Given that the system reaches quasi-static chemical equilibrium relative to the mechanical timescales, the closed bond density *ρ*(*r*) is determined by the balance between the constant association rate *k*_on_ and the force-dependent dissociation rate *k*_off_ (*F*_*b*_). The explicit analytical derivation for *ρ* is presented in S2.3.

#### 2.1.4 Adhesion Chemical Energy (*U*_adh_)

The formation of ligand-receptor bonds leads to a reduction in chemical energy, *U*_adh_, expressed as:

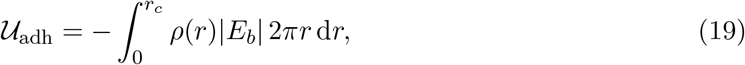

where 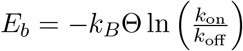 represents the binding free energy per bond.

The initial mechanical state is identified by minimizing the total free energy *U* (*R, h*). However, optimization is constrained by the conservation of volume and ion content, which couples *R* and *h* via *f* (*R, h*) = 0 (derived in section S2.1). Thus, we solve:

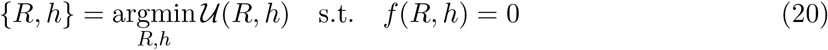

### 2.2 Dynamic cell evolution under osmotic shock

#### 2.2.1 Water and ion balance equation

Based on the principles of water and ion mass balance (44), the transmembrane water flux *J*_*w*_ is driven by the hydrostatic (Δ*P*) and osmotic (ΔΠ) pressure gradients:

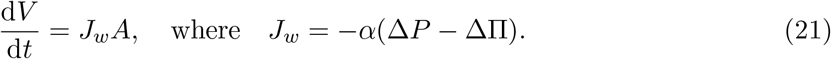

where *α* is the hydraulic permeability. The hydrostatic pressure follows the Young-Laplace law Δ*P* = 2*σ*_cort_*d/R*, dependent on the cortical stress *σ*_cort_ and thickness *d* (45). The osmotic pressure difference is ΔΠ = Π_in_ *−* Π_out_, where the osmotic pressures obey the Van’t Hoff relation Π = (*n/V*)*R*_*k*_Θ (46, 47), with *R*_*k*_ representing the gas constant. Coupled to volume changes, intracellular osmolarity is regulated by the net ion flux *J*_*n*_, which comprises active pumping (*J*_*a*_) and passive leakage (*J*_*p*_):

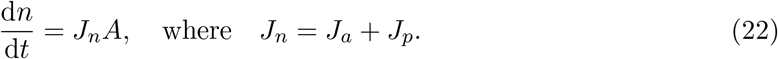

Active transport acts as a homeostatic feedback mechanism, driven by the ion pump pressure ΔΠ_*c*_ (29):

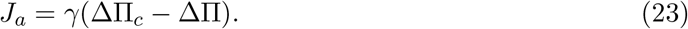

Passive transport *J*_*p*_ is mediated by mechanosensitive (MS) channels responding to the cortical stress *σ*_cort_. We model this flux as a piecewise function:

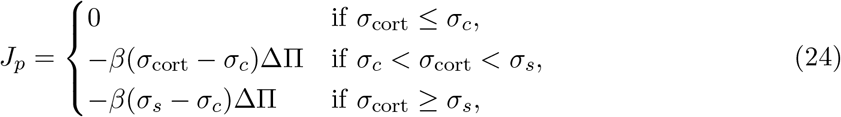

where *β* determines the channel sensitivity. The system is characterized by a threshold stress *σ*_*c*_ for channel opening and a saturation stress *σ*_*s*_, beyond which the channels are fully open.

#### 2.2.2 Power balance equation

To describe the dynamic evolution of the cell under osmotic shock, we enforce a global thermodynamic power balance, postulating that the system is primarily driven by the rate of work done by the hydrostatic pressure difference:

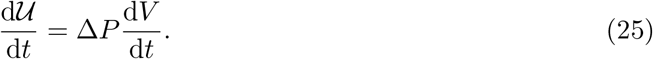

Given that the mechanical work is mainly dissipated through cortical deformation, this formulation implies that variations in other energy components (*U*_SF_ + *U*_sub_ + *U*_adh_) are negligible during the rapid osmotic shock (see section S2.2 for the detailed derivation). This aligns with the biophysical observation that focal adhesion remodeling occurs on a much slower timescale than osmotic response.

### 2.3 Hill-type Active Strain Coupling

Cellular mechanical morphology is inherently dynamic, evolving during processes such as spreading, migration, contraction, and adaptation to osmotic changes. This necessitates a modeling framework capable of capturing such dynamic mechanics. In this context, the molecular clutch framework has emerged as a leading paradigm for elucidating cell mechanosensitivity and has been shown to reproduce key features of cell spreading and migration (31, 33, 48, 49). It provides a fundamental link between macroscopic morphological changes and microscopic molecular events, serving as a basis for predicting cellular responses to mechanical cues. Cell spreading is driven by the competition between actin polymerization (*v*_*p*_) and retrograde flow (*v*_flow_). The process shifts from an initial expansion (*v*_*p*_ *> v*_flow_) to a contraction phase (*v*_flow_ *> v*_*p*_), finally reaching a homeostatic state where *v*_*p*_ ≈ *v*_flow_.

However, standard implementations of these models are generally confined to quasi-static or steady-state assumptions. While effective for quasi-static processes evolving over timescales exceeding 100 s, they may overlook transient dynamics induced by rapid physiological stimuli. Examples include pulsatile shear stress from heartbeats, cyclic stretch during respiration, and acute osmotic shocks—events that operate on timescales distinct from slow cell spreading and migration. In these fast-evolving scenarios, the instantaneous kinetic response of traction forces cannot be effectively captured by existing static frameworks. Moreover, although we established the global power balance in Eq. (25), that formulation does not yet explicitly account for the dynamic response of the traction force *T*. To address this limitation, we introduce a dynamic Hill-type law that couples the active cytoskeleton strain *ε*_a_(*t*) to the cell edge velocity (see Fig. 3).

**Figure 3:**
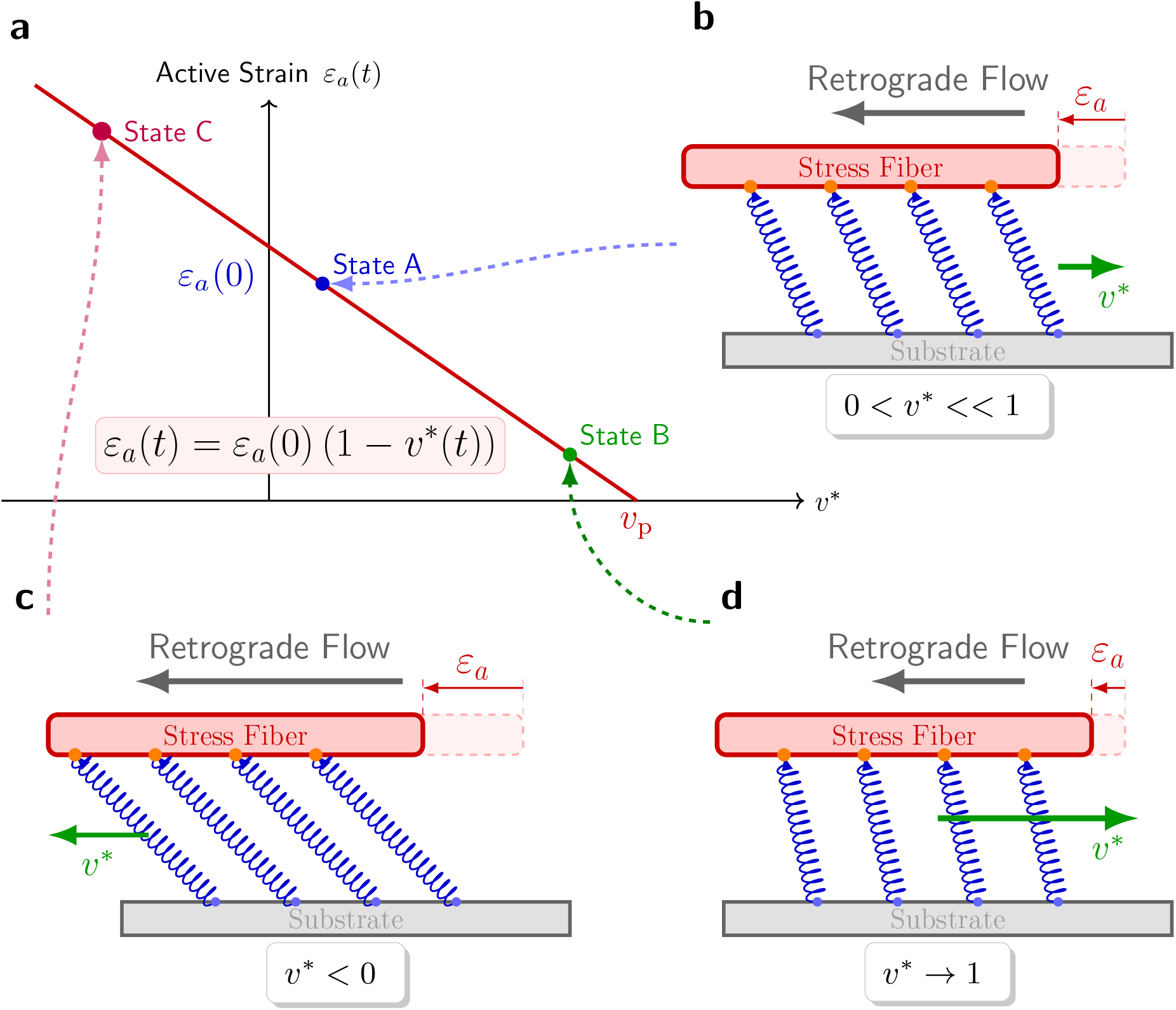
Schematic of the dynamic constitutive relationship for active strain *ε*_a_. (a) Profile of active strain *ε*_a_ as a function of the normalized edge velocity *v*\*. (b) Low velocity regime: Slow expansion results in negligible kinematic decay of *ε*_a_. (c) Contractile regime: Inward motion (negative *v*\*) drives the amplification of active strain (strain growth). (d) High velocity regime: Rapid expansion leads to significant kinematic decay, suppressing the active strain.

Based on the motor-clutch model, the cell edge velocity 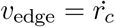 is defined as the net result of the constant actin polymerization rate *v*_p_ and the retrograde flow velocity *v*_flow_:

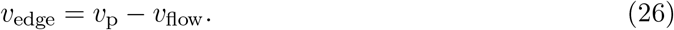

While recent studies have highlighted the complex coupling between retrograde flow and stress fiber contractility via molecular clutch dynamics (49) or stretch (50), we propose a simplified linear ansatz to capture this behavior at the continuum level:

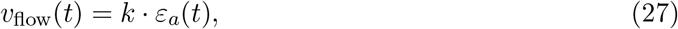

where *k* is a kinetic constant. We determine the constant *k* by identifying the stall condition (where *v*_edge_ = 0 implies *v*_flow_ = *v*_p_ and *ε*_*a*_ = *ε*_*a*_(0)). This leads to the constitutive relation:

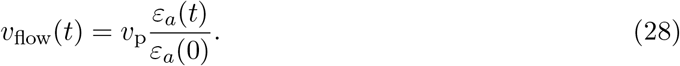

Combining Eq. (28) with Eq. (26) yields the expression *v*_edge_(*t*) = *v*_p_[1 *− ε*_*a*_(*t*)*/ε*_*a*_(0)]. By defining the dimensionless velocity *v*\* = *v*_edge_*/v*_p_ and rearranging terms, we arrive at the relation for active strain dynamics:

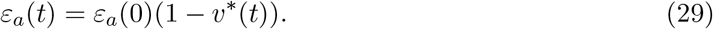

Figure 3 schematically illustrates the constitutive relationship proposed in Eq. (29). As shown in Figure 3a, the active strain *ε*_*a*_(*t*) manifests as a linearly decreasing function of the edge velocity *v*_edge_. To ensure the physical consistency of inward traction forces, we introduce a cut-off condition where *ε*_*a*_(*t*) becomes zero when *v*_edge_ *≥ v*_p_, effectively preventing any unphysical force reversal.

Specifically, three characteristic dynamic regimes (State A, B, C) are identified (Fig. 3b-d): State A Steady Spreading: Under conditions of moderate edge protrusion (0 *< v*\* *≪* 1), such as typical cell spreading, the active strain is maintained at an intermediate level (Fig. 3b). State B (Rapid Expansion): Conversely, during rapid edge advancement (e.g., hypotonic shock), the edge velocity approaches its maximum limit (*v*\* *→* 1). This minimizes the relative retrograde flow, causing *ε*_*a*_(*t*) to decrease drastically (Fig. 3d). State C Retraction: In contractile scenarios (e.g., hypertonic shock), the edge velocity becomes negative (*v*\* *<* 0). This inward motion leads to an increase in *ε*_*a*_(*t*) (Fig. 3c). Substituting the dynamic active strain equation from Eq. (29) into the quasi-static solution Eq. (14) yields a rate-dependent displacement field 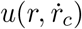. Consequently, the power balance in Eq. (25) should be reformulated to obtain the dynamic solution.

To efficiently solve this coupled system, we developed a semi-analytical strategy that reduces the two-dimensional geometry search to a one-dimensional root-finding problem via geometric substitution, solving the radius from an explicit quadratic mapping at each time step while enforcing the global power balance through an outer iterative loop (see sections S2.6 and S2.7 for the full derivation and implementation flowchart).

#### 2.3.1 Dynamic Traction Force Derivation

To physically analyze the kinematic effect on the traction force *T*, we specifically focus on the edge traction *T*_edge_. By evaluating Eq. 14 at the cell edge (*r* = *r*_*c*_), we obtain the analytical expression for the edge displacement *u*(*r*_*c*_):

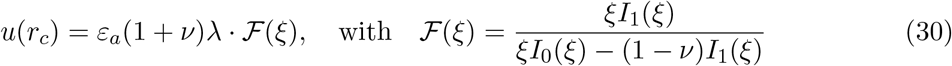

where *ξ* = *r*_*c*_*/λ* denotes the dimensionless adhesion radius and *λ* is the screening length.By incorporating the kinematic effect from Eqs. (29) and (30) into the traction force definition (Eq. (8)), we derive the dynamic cell edge traction force *T*_edge_:

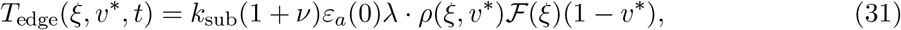

for which a detailed derivation can be found in section S2.3. While an exact solution for the temporal evolution of *F*(*ξ*(*t*)) is not available, we can capture the system’s essential behavior through asymptotic analysis of the geometric factor *F*(*ξ*) in two limiting regimes: small adhesion (*ξ →* 0) and large adhesion (*ξ →* ∞).

#### 2.3.2 Asymptotic Analysis of Steady-State Traction Force

To establish a theoretical baseline, we examine the asymptotic limits of the static solution. In the small adhesion limit (*ξ →−* 0), Taylor expansions (*I*_0_ ≈ 1, *I*_1_ ≈ *ξ/*2) simplify the geometric factor to *F* ≈ *ξ/*(1 + *ν*), yielding a linear displacement *u*(*r*_*c*_) ≈ *ε*_*a*_*λξ*. By substituting the asymptotic bond density *ρ*(*ξ*) ≈ *ρ*_0_ *−* (*C*_0_*ε*_*a*_*λ*)*ξ* derived in section S2.4, the edge traction becomes:

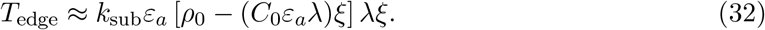

Neglecting the higher-order quadratic term *O*(*ξ*^2^), this reveals a geometric strengthening regime where traction scales linearly with radius (*T*_edge_ ∝ *ξ*).

Conversely, as *ξ →* ∞, the ratio *I*_1_*/I*_0_ converges to unity, stabilizing both displacement (*u*(*r*_*c*_) ≈ *ε*_*a*_(1 + *ν*)*λ*) and bond density (*ρ → ρ*_∞_, see section S2.4). Consequently, the traction saturates to:

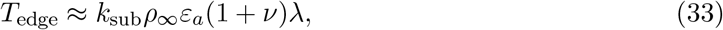

indicating that geometric expansion yields negligible gain in traction transmission at this limit. Physically, this implies that the edge traction is determined solely by the local mechanical environment within a screening length *λ*, rendering the remote central region mechanically irrelevant.

#### 2.3.3 Asymptotic analysis of dynamic traction force

Building on the static analysis above, we consider the dynamic response where *T*_edge_ varies with the evolving strain *ε*_*a*_(*t*). In this section, we analyze the rate of traction change 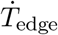 in the asymptotic limits of small and large *ξ*.

##### The geometric-kinematic competition regime (small *ξ*)

In the limit of small adhesion radii (*ξ →* 0), we incorporate time dependence into Eq. (32):

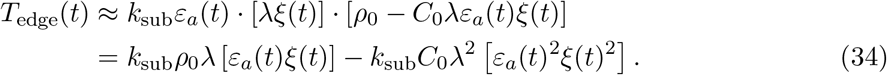

Substituting the dynamic active strain *ε*_*a*_(*t*), we group the constants into linear (*C*_1_) and quadratic (*C*_2_) coefficients:

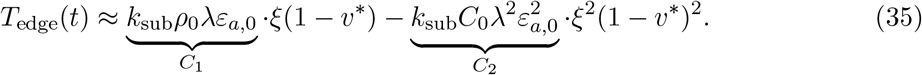

To determine the evolution rate 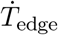, we differentiate with respect to time using the chain rule. Let Ψ(*t*) = *ξ*(*t*)[1 *− v*\*(*t*)]. The time derivative is 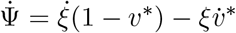. The traction force rate becomes:

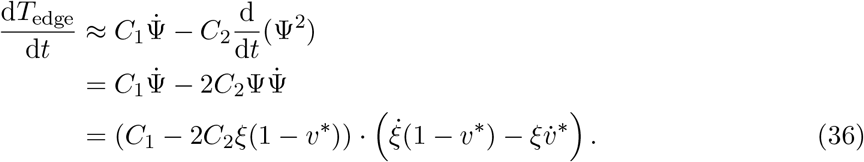

In the limit of *ξ*, 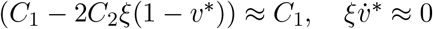, then Eq. (36) is simplified to:

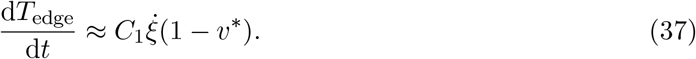

This equation reveals a fundamental geometric-kinematic competition in traction dynamics. Specifically, in the low *v*\* regime (typical of physiological cell spreading), the process is geometrically dominated: the expansion of adhesion radius 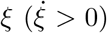 drives a net accumulation of traction force, as a larger *ξ* induces larger displacement *u*(*r*_*c*_) (see Eq. (32)), significantly out-weighing the negligible contribution from velocity decay. Conversely, in the high-velocity limit (*v*\* *→* 1), the system shifts to a kinematically dominated regime. Here, traction force decays because the rapid edge velocity impedes the buildup of active strain *ε*_*a*_(*t*)—and consequently displacement *u*—as described by the Hill-type coupling in Eq. (29).

##### The kinematically-dominated regime (large *ξ*)

In the asymptotic limit of large adhesion radii (*ξ →* ∞), consistent with the steady-state solution in Eq. (33), the cell edge traction force *T*_edge_ enters a saturation phase. Here, the edge integrin density *ρ*(*r*_*c*_) converges to a stable plateau *ρ*_∞_, while the edge displacement *u*(*r*_*c*_) approaches the limit *u*(*r*_c_) ≈ *ε*_*a*_(*t*)(1 + *ν*)*λ*. Consequently, *T*_edge_ decouples from the instantaneous radius *r*_*c*_, becoming driven solely by the time-varying active strain:

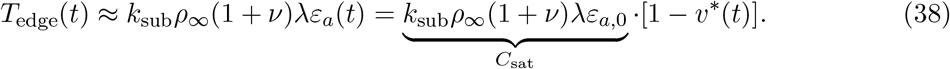

Differentiating with respect to time yields a simplified relation:

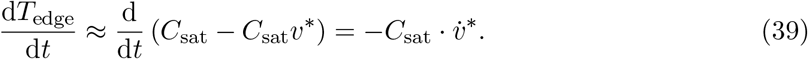

In this saturated regime, the explicit dependence on geometric growth 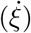 vanishes because both the displacement field and bond density have reached their asymptotic limits. The traction evolution is thus entirely dictated by the acceleration of spreading. Consequently, any increase in spreading velocity 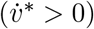 directly translates to a decrease in traction force 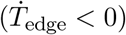.

#### 2.3.4 Illustrations

To quantify the geometric-kinematic competition, we simulate the evolution of *ξ*(*t*) and normalized traction 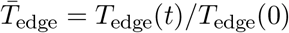 driven by a prescribed velocity field *v*\*(*t*). The normalized radius *ξ*(*t*) evolves cumulatively via kinematic integration:

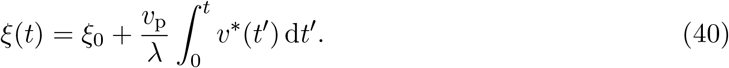

Substituting *ξ*(*t*) into the traction solution yields the dynamic response. We employ a representative velocity profile comprising acceleration and deceleration phases to capture the full spreading spectrum (Fig. 4a). As spreading proceeds, *ξ*(*t*) increases monotonically from distinct initial values (*ξ*_0_ = 1, 2, 5, Fig. 4b). As predicted by our asymptotic analysis in section 2.3.3, the evolution of normalized traction force 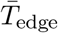 exhibits a fundamental divergence depending on the initial cell size *ξ*_0_ (Fig. 4c). Specifically, for the small adhesion case (*ξ*_0_ = 1, blue line), the traction force rises monotonically throughout the process. This confirms the prediction of the geometrically-dominated regime, where the geometric strengthening effect (in Eq. (37)) overwhelms the decay effect induced by velocity acceleration. In sharp contrast, large cells (*ξ*_0_ = 2, 5, red/green Lines) display a non-monotonic response, characterized by a distinct dip during the acceleration phase (*t* ≈ 4 *−* 7s). This behavior is consistent with the kinematically dominated regime, where geometric saturation renders the cell mechanics highly sensitive to the rate of velocity change *v*?*. As derived in Eq. (39), the positive acceleration 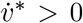 directly drives a negative time derivative 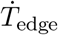, thereby causing the observed transient decline in traction force.

**Figure 4:**
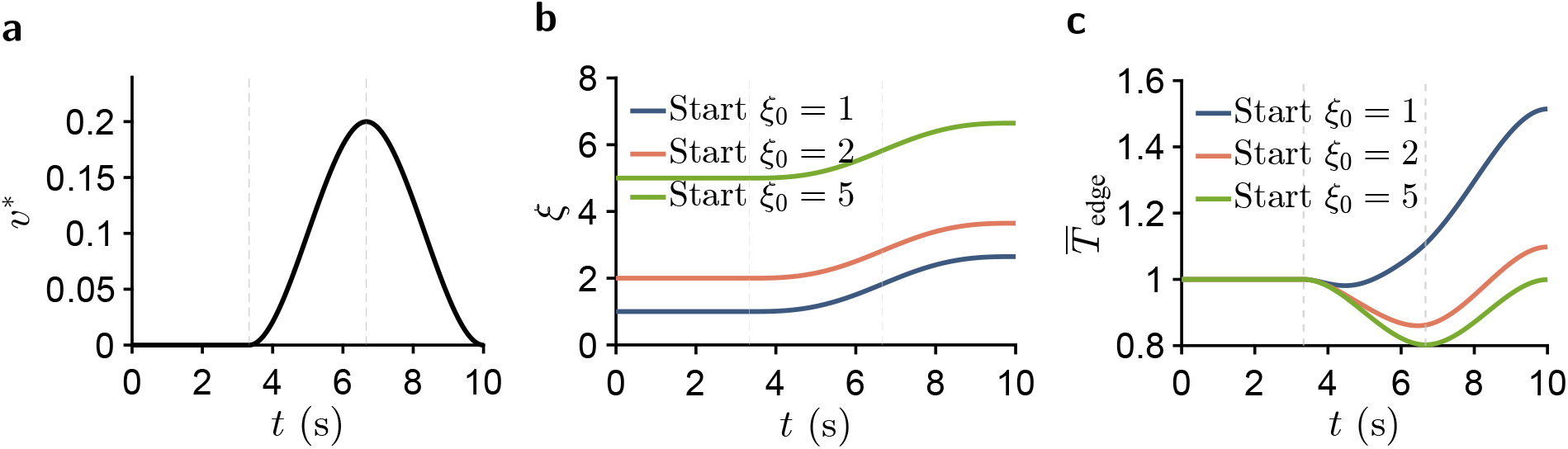
Size-dependent traction force divergence under dynamic loading. (a) The prescribed time-dependent velocity field *v*\*(*t*) imposed, characterized by distinct acceleration and deceleration phases. (b) The resulting temporal evolution of dimensionless cell size *ξ*(*t*) for three distinct initial sizes (*ξ*_0_ = 1, 2, 5). (c) The corresponding evolution of normalized traction force *T* _edge_. The response depends on the initial size *ξ*_0_: small adhesion (*ξ*_0_ = 1, blue) shows a monotonic force increase (Geometric-Dominated), whereas large cells (*ξ*_0_ = 5, green) display a transient force dip (Kinematically-Dominated) during the rapid expansion phase.

To isolate geometric contributions from kinematic fluctuations, we simulated idealized constant-velocity scenarios 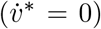 representing slow spreading (*v*\* = 0.1), contraction (*v*\* = ™0.05), and steady states (*v*\* = 0) in Fig. S3a. Under these conditions, the acceleration term in Eq. (36) vanishes, implying that traction dynamics are dominated by the geometric contribution in this control limit. This prediction is consistent with our results: unlike the dynamic cases, the normalized traction force 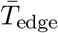 exhibits strictly monotonic behavior with respect to *ξ* (Fig. S3b–d). Thus, the model captures traction evolution under quasi-static controls spanning both spreading and detachment phases. Moreover, this control simulation supports the interpretation that the non-monotonic dip observed in Fig. 4 reflects the kinematic contribution driven by dynamic acceleration.

## 3 Results and Discussion

### 3.1 Osmotic Response and Stiffness-Associated Recovery Divergence

#### 3.1.1 Cellular Dynamics under Hypotonic Shock

We initially employed the model, parameterized according to (table S2), to simulate the response of adherent cells to hypotonic shock. As illustrated in Fig 5(a-c), the variables 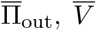, and 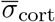 represent values normalized to their initial states at *t* = 0 s. Upon reducing the normalized external osmotic pressure 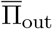 to one-third of its baseline (Fig 5a), the cell volume 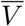 expands drastically (Fig 5b), concurrent with a sharp increase in the sphere cap area 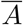 (Fig S4b) and radius 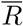 (Fig S4c). Through Eq. (4), these geometric expansions induce a transient increase in the cortical stress 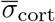 (Fig 5c). Nevertheless, once the osmotic perturbation ceases, 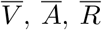 and 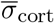 demonstrate a recovery behavior (Fig. 5(b-c), Fig. S4(b-c)). These observations are consistent with our intuitive physical expectations.

**Figure 5:**
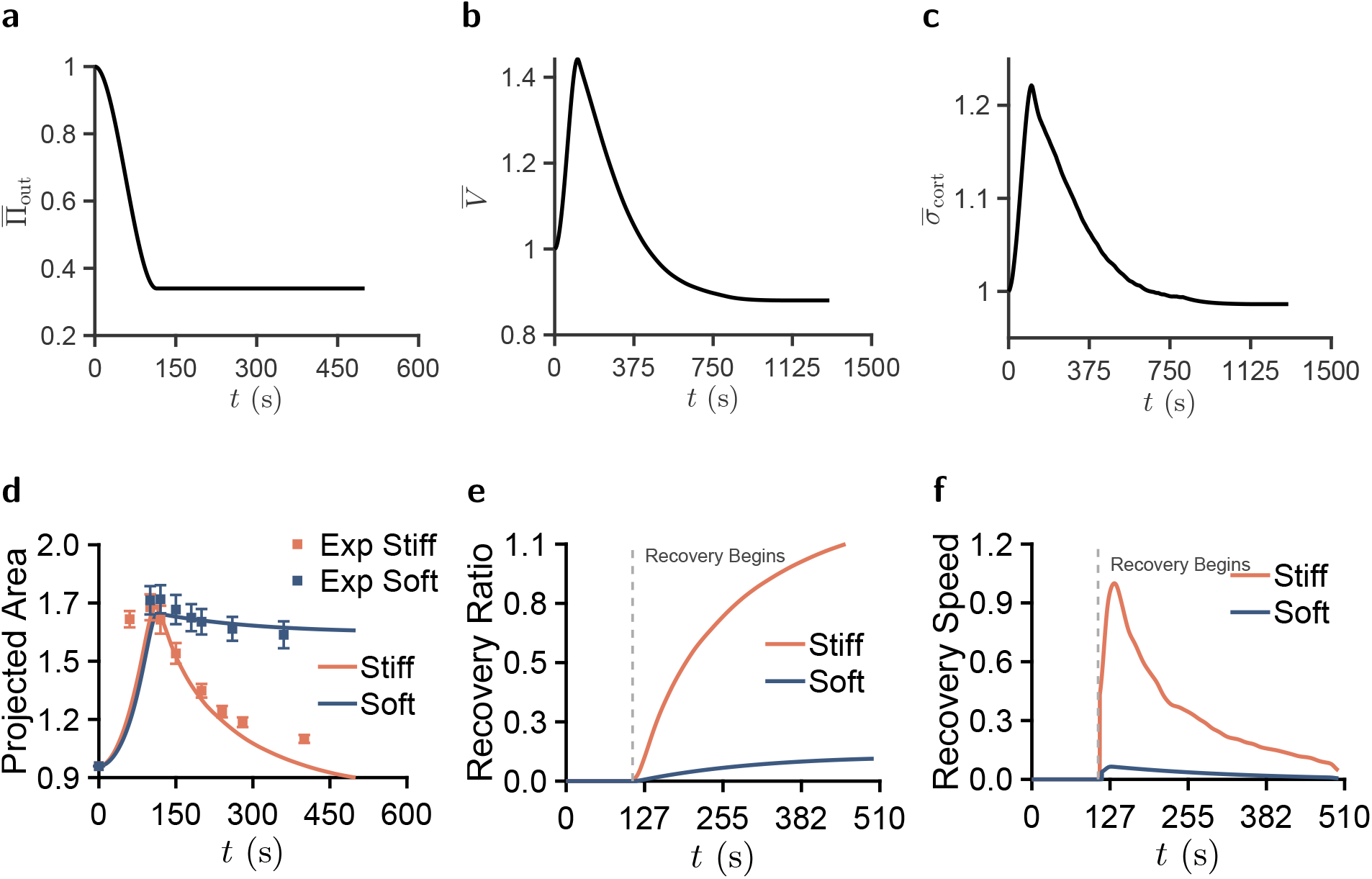
Dynamic response of adherent cells to hypotonic shock. All variables are normalized to their initial values (*t* = 0), with experimental data used for model-experiment comparison of stiffness-associated recovery dynamics. (a) The temporal profile of the external osmotic pressure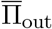, which is stepped down to 1*/*3 of the baseline to mimic hypotonic shock. (b) Time evolution of the normalized cell volume 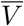. The cell undergoes rapid expansion followed by a recovery phase. (c) Time evolution of the normalized cortical stress 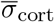, showing an instantaneous surge due to cell swelling and subsequent relaxation. (d) Temporal evolution of the normalized projected area. The solid lines and points represent the model simulation and experimental data. (e) Recovery ratio of stiff cells and soft cells. (f) Recovery speed of stiff cells and soft cells.

Because the simulated osmotic response involves coupled changes in cell volume, radius, and cortical stress, we further examined whether the predicted recovery behavior is sensitive to elastic parameters associated with volume–shape coupling. In section section S3.2, we varied the cortical Poisson ratio *ν*_*m*_ and the stress-fiber Poisson ratio *ν*_SF_ (section S3.2). The results show that changing *ν*_*m*_ affects the normalized radius response, whereas the normalized stress and edge traction remain only weakly affected by the tested Poisson-ratio variations.

Pathological transformations are frequently accompanied by significant mechanical alterations, exemplified by the distinct cortical softening observed in metastatic MDA-MB-231 cells relative to non-tumorigenic MCF-10A cells (51). Motivated by this mechanical contrast, we simulated hypotonic shock (Fig. 5a) using different cortical Young’s moduli measured by AFM indentation (Fig. S4a, *p <* 0.01; experimental details in section S1). As illustrated by the red line (stiff cells) and blue line (soft cells) in Fig. 5d, the normalized projected area (defined in Fig. S1) revealed a stiffness-associated divergence: while both cell types exhibit projected area recovery, the stiffer parameter set recovers more rapidly than the softer one.

We compared this simulation with a previously reported MCF-10A and MDA-MB-231 projected-area dataset (52), in which both cell types were subjected to the same hypotonic protocol (start from t=0 s)300 mOsm *→* 100 mOsm) while projected area was monitored by microscopy. The experimental trends (symbols in Fig. 5d) are consistent with the model: the stiffer MCF-10A cells (*n* = 14) recover faster than the softer MDA-MB-231 cells (*n* = 15). This cell-line comparison supports an association between cortical mechanical state and recovery kinetics, while other cell-line-dependent properties may also contribute.

To further assess this interpretation, we refer to the previously reported actin-perturbation experiments (52). In that work, Cytochalasin D treatment of MCF-10A cells disrupted actin polymerization, reduced cortical stiffness, and slowed hypotonic projected-area recovery within the same cell type. Conversely, Jasplakinolide treatment, which stabilizes F-actin and promotes restoration of the cortical mechanical state, accelerated projected-area recovery. These opposing perturbations support cortical mechanical state as an important contributor to osmotic recovery in the tested system. Ion transport, adhesion remodeling, and other biochemical regulators may also contribute to the measured recovery kinetics.

We therefore treated cortical stiffness as a measurable descriptor of the broader actin-cortex mechanical state and quantified the associated recovery differences using the normalized recovery fraction, 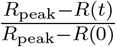, where *R*_peak_ is the cell radius at maximum expansion and *R*(0) is the initial radius, together with the recovery speed 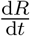. As illustrated in Fig. 5(e-f), the stiffer condition exhibits both a higher recovery extent and faster recovery kinetics than the softer condition.

#### 3.1.2 Stiffness-Associated Recovery Divergence

To elucidate the physical origin of this stiffness-associated divergence, we analyze the transmembrane water flux *J*_*w*_ (Eq. (21)), which is driven by the interplay between hydrostatic (Δ*P*) and osmotic (ΔΠ) pressure differences. Upon hypotonic shock, the sharp osmotic difference ΔΠ surge (Fig. 6a) acts as the primary driving force for water influx. Initially, Δ*P* drops (in Fig. 6b), becoming insufficient to counterbalance ΔΠ. This hydrodynamic imbalance (*J*_*w*_ ∝ ΔΠ *−* Δ*P >* 0) drives the rapid swelling phase (Fig. 6c). As ion transport mechanisms engage and elastic recoil takes effect, the cell gradually recovers towards a new steady state where Δ*P* ≈ ΔΠ. While stiffness-associated divergence is minimal during the initial swelling phase (Fig. 5d)—where dynamics are dominated by the large osmotic gradient—it becomes a distinctive characteristic of the recovery phase. Consequently, the following analysis focuses on the physical origins of this stiffness-associated recovery. According to Fig. 6b, the divergence in recovery behavior arises from the cell’s capability to generate a hydrostatic pressure recovery. Stiff cells exhibit a distinct pressure rebound, whereas soft cells settle into a low-pressure steady-state. To explain this, we decomposed the pressure rate d(Δ*P*)*/*d*t*:

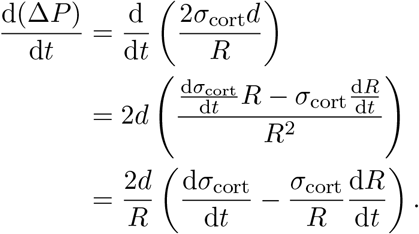

We designate the terms 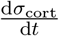 and 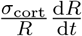 as the stress rebound and geometric rebound terms, respectively. As illustrated in Fig. 6e, stiff cells are characterized by a deep negative trough in the geometric rebound term, driven by rapid radius contraction (*dR/dt <* 0, in Fig. S5a). In the rate equation, this negative geometric rebound term functions as the primary driver for pressure recovery (Fig. 6f), far exceeding the contribution from the stress rebound term (Fig. 6d, Fig. S5b). Conversely, soft cells generate only minimal magnitudes for both terms. These results suggest that higher stiffness facilitates faster geometric recoil, which amplifies hydrostatic pressure recovery and promotes water efflux. Such mechanical recoil provides one contribution to osmotic adaptation, alongside cell-type-dependent RVD pathways regulated by ion channels and transporters. The cell cortex acts like a stiff spring: the stronger recoil force drives rapid contraction, instantly boosting the hydrostatic pressure.

**Figure 6:**
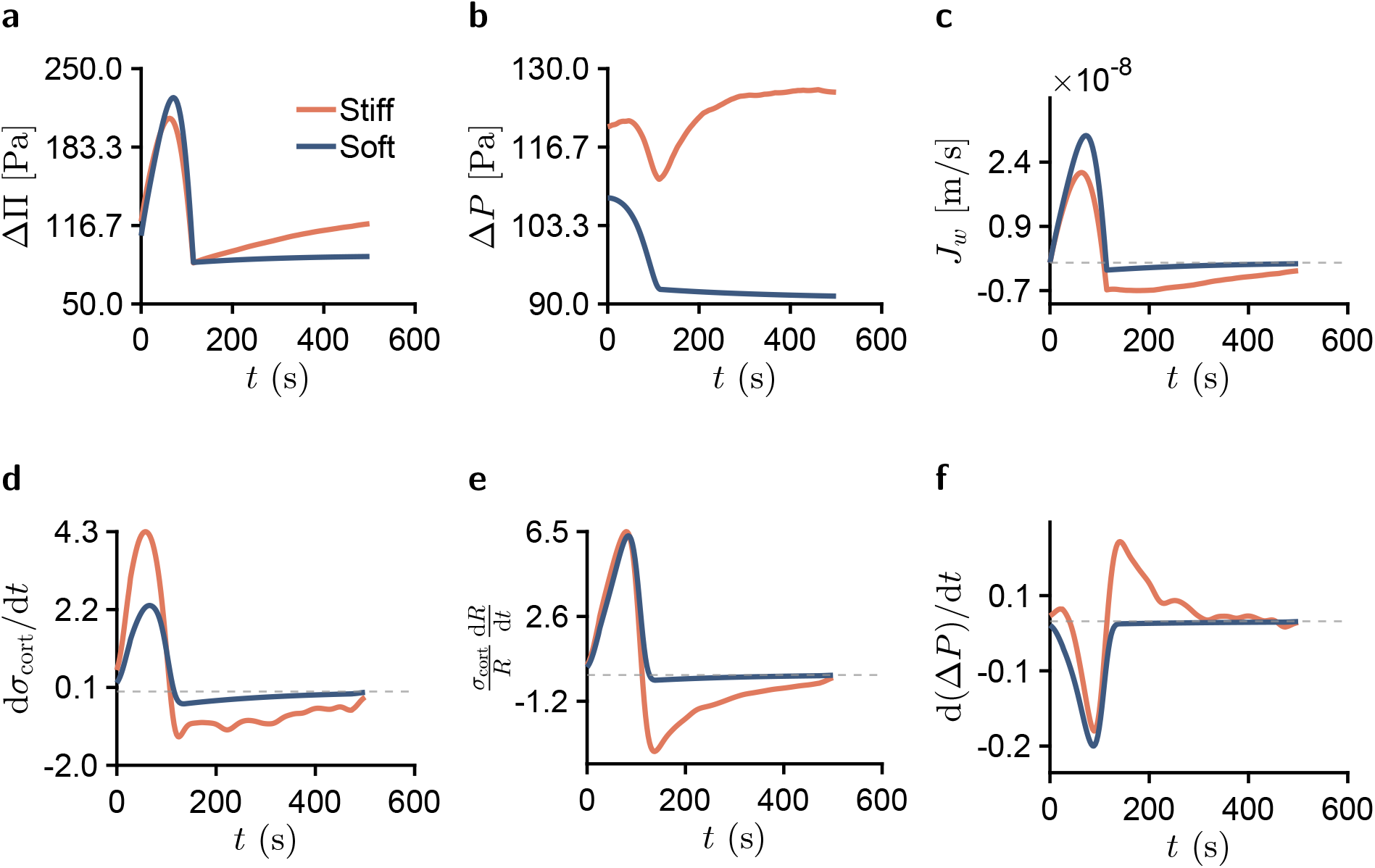
Hydrodynamic balance and its mechanical origin. (a-c) Macroscopic Dynamics: The osmotic gradient ΔΠ (a) induces a hydrostatic pressure Δ*P* (b). The interplay between these pressures dictates the water flux *J*_*w*_ (c), where stiff cells show an earlier reversal from influx to efflux. (d-f) Microscopic mechanism: Decomposition of the pressure rate d(Δ*P*)*/*d*t* (f) reveals that stress accumulation (d) overwhelms geometric relaxation (e).

### 3.2 Competition between Geometric Strengthening and Kinematic Decay

#### 3.2.1 The Kinematically-Dominated Regime: Transient Force Decay

Under osmotic shock, cell morphology undergoes drastic alterations, driving dynamic variations in traction forces. Fig. 7a illustrates the evolution of the cell throughout this process (15). The TFM heatmaps (top panels) reveal a substantial decay of traction stress during the swelling phase, evidenced by the shift from high-stress to low-stress regions, followed by a transient recovery and subsequent decay during the shrinking phase. Concurrently, morphological tracking (bottom panels) confirms rapid cell swelling (0 – 150 s) prior to the onset of shrinkage. Crucially, this observed decline in traction force during area expansion stands in sharp contrast to conventional spreading mechanical theories (42), which postulate a monotonic scaling between traction and adhesion size. This discrepancy underscores the inadequacy of static frameworks in capturing such transient dynamic effects.

**Figure 7:**
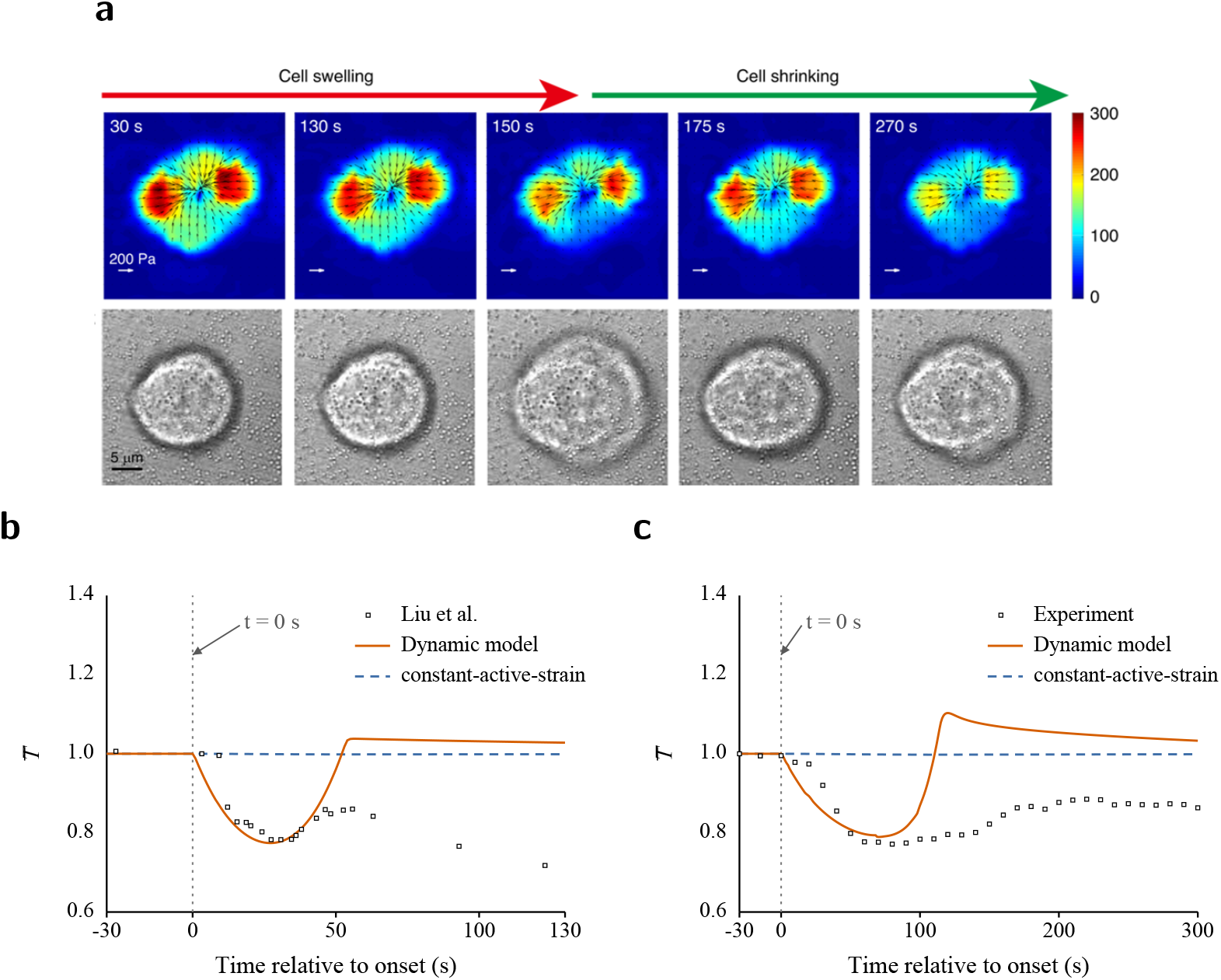
Model–experiment comparison of traction dynamics. (a) Traction-stress heatmaps and cell morphology reproduced from Liu et al. (15), showing the traction response during hypotonic swelling and recovery. (b,c) Temporal evolution of the normalized edge traction force, 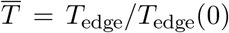. Time was aligned by setting *t* = 0 at the onset of the projected-area response, and curves were normalized by the pre-shock value. Lines denote the dynamic model and the constant-active-strain model; symbols denote experimental data. Panel (b) shows slightly adherent C2C12 cells from Liu et al. (15) (start from t=0 s, hypotonic protocol (300 mOsm *→* 200 mOsm),*n* = 5),substrate Young’s modulus=2.5kPa), and panel (c) shows our MCF-10A (start from t=0 s hypotonic protocol (300 mOsm *→* 100 mOsm), *n* = 1, subatrate Young’s modulus=10kPa).

This traction decrease can be understood from the asymptotic analysis in section 2.3.3, which gives a kinematically dominated regime for large cells. In this regime, the dynamic constitutive law predicts that the kinematic decay effect (which reduces stress) can exceed the geometric strengthening effect (which increases stress). Consequently, a rapid hypotonic expansion is expected to trigger a transient decrease in traction force, providing a mechanics-based explanation for the experimentally observed force decrease.

We next compared the model-predicted traction response with experimental data from our TFM measurements and the dataset reported by Liu et al. (15). As shown in Fig. 7(b,c), both datasets exhibit a rapid decrease in normalized edge traction during the initial hypotonic shock, reaching approximately 0.8 of the pre-shock value during rapid swelling.

The dynamic model reproduced this initial traction-loss trend, whereas the constant-active-strain model remained close to the pre-shock baseline. This contrast suggests that, after the adhesion geometry has adjusted, the geometric contribution alone has a limited dynamic range, and the velocity-dependent active-strain response provides the additional kinematic contribution needed to capture the rapid traction decrease.

The late response follows a different trajectory from the rapid-swelling phase. The model is consistent with the initial traction-force decrease during hypotonic swelling, whereas the (15) trace continues to decline and our experimental trace shows incomplete recovery. These differing late-stage trajectories suggest that traction recovery after the initial osmotic expansion involves additional longer-timescale cellular regulation. Possible contributors include actin-cortex remodeling, adhesion turnover, partial weakening of integrin-mediated adhesion, regulatory volume recovery, and ion-transport-mediated adaptation. These processes can alter the effective traction-generating state of the cell after the early swelling response and may therefore influence whether traction returns to its pre-shock level.

#### 3.2.2 Transition Mechanism: From Decay to Strengthening

After this model-experiment comparison, we extend the analysis to a broader parameter space. According to Eq. (39), in the large *ξ* regime, the cell edge traction *T*_edge_ becomes kinematically dominated and exhibits an inverse dependence on velocity *v*\*. Specifically, traction decays during fast expansion but intensifies during rapid contraction, a behavior consistent with experimental findings (15). Conversely, Eq. (37) suggests the existence of a geometrically-dominated regime. For instance, during the initial phase of cell spreading onto the substrate, traction force increases due to the expanding adhesion area. Similarly, during the phase of cell detachment from the substrate, the reduction in adhesion area leads to a concomitant decrease in traction force.

Synthesizing these observations, we predict that the interplay between *v*\* and *ξ* yields four distinct dynamic phases of traction evolution. To evaluate this prediction, the edge traction *T*_edge_(*ξ, v*\*, *t*) is computed via Eq. (31). For a given initial state *ξ*(0) and instantaneous velocity *v*\*, we examine the traction change after a time interval Δ*t*, where the time-evolution of *ξ*(*t*) is obtained from Eq. (40). We define the normalized traction variation as 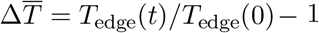. This metric serves as an indicator to determine whether the kinematic process *v*\* induces strengthening 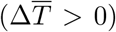 or decay 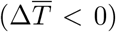. By systematically computing this metric across the state space spanned by *ξ* and *v*\*, we construct the schematic mechanism and phase diagram shown in Fig. 8.

**Figure 8:**
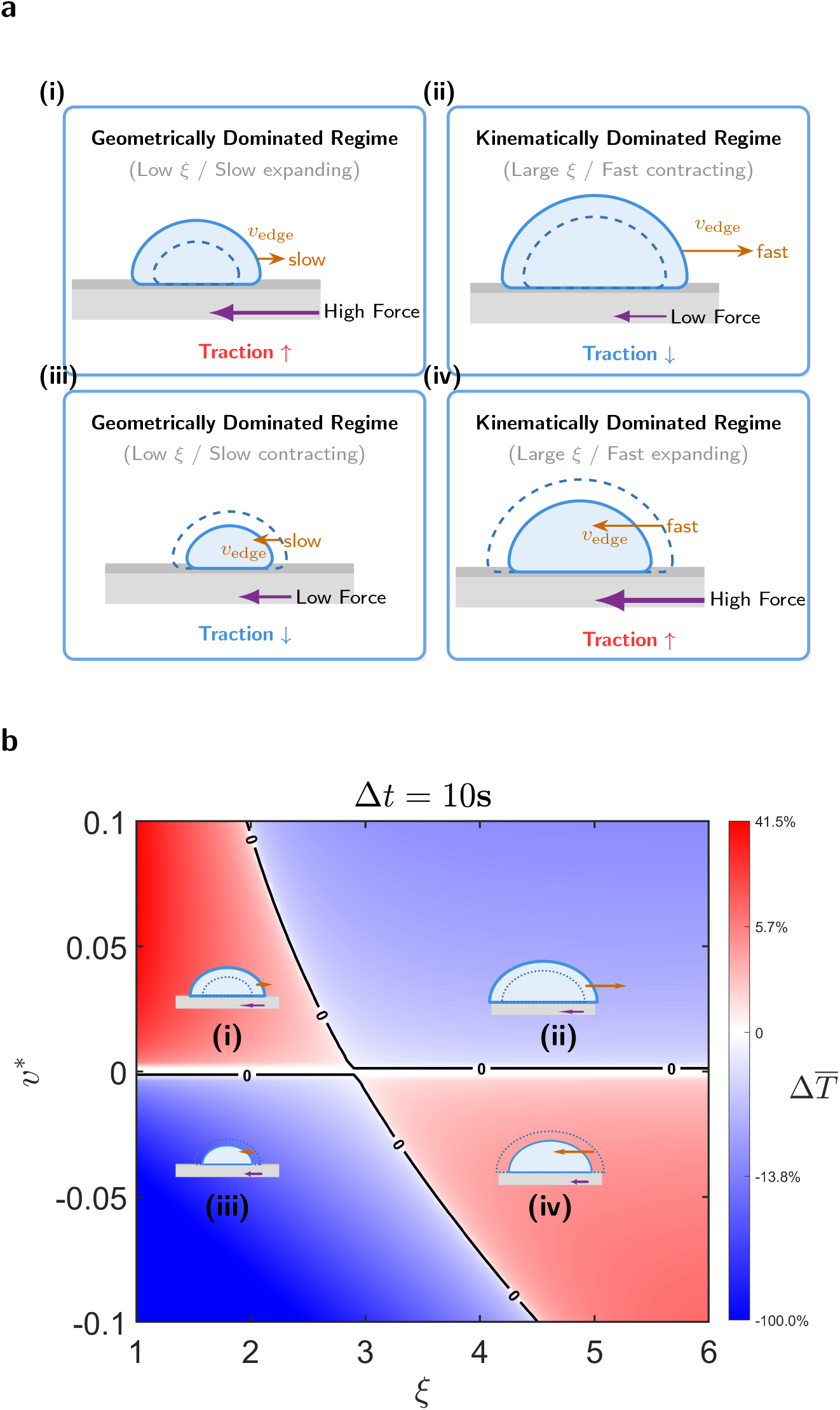
Mechanism and phase map of the geometric-kinematic competition. (a) Schematic illustrating the opposing roles of geometric and kinematic effects. (b) Phase diagram of traction force at an observation time of Δ*t* = 10 s. The solid line denotes the zero-isocline (Δ*T* = 0), partitioning the phase space into two distinct domains: a strengthening zone (red) and a weakening zone (blue).

The geometric-kinematic competition, conceptually illustrated in Fig. 8a, is mapped in the phase diagram of Fig. 8b. This color map partitions the system’s response into two distinct physical regimes separated by the zero-isocline (defined by Δ*T* = 0): a decay zone (blue) and a strengthening zone (red).

##### Geometrically dominated Regime

Panels (i) and (iii) illustrate the geometrically-dominated regime, characterized by a small adhesion size *ξ* and low velocity *v*\*. As demonstrated by the asymptotic analysis in section 2.3.2 and Eq. (37), the system aligns with mechanical intuition: adhesion expansion generates higher traction force, while shrinkage leads to relaxation. This behavior physically corresponds to the initial stages of cell spreading or cell detachment (53). Notably, Fig. 8b illustrates the kinematic influence on the phase diagram: a faster *v*\* corresponds to regions of higher color intensity, indicating a greater magnitude of force change 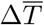. Consistent with Eq. (37), the traction rate scales linearly with the geometric expansion rate 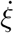, which is directly proportional to *v*\*. Thus, a higher velocity acts as a multiplier: it increases the rate of force accumulation during expansion (deep red) or accelerates force relaxation during contraction (deep blue), leading to the observed saturation in color depth.

##### Kinematically dominated Regime

Panels (ii) and (iv) illustrate the kinematically dominated regime, typically observed at large adhesion size where the geometric sensitivity saturates. In these regimes, the traction evolution is shaped primarily by the velocity-dependent active strain, *ε*_*a*_(*v*\*), as defined by the Hill-type constitutive law (Eq. (29)). In the case of rapid expansion (Case ii, *v*\* *>* 0) : The positive velocity reduces active contractility (*ε*_*a*_ ↓, Fig. 3d), which directly translates to a net decrease in traction force according to Eq. (39) (as depicted in Fig. 8a (ii)). This mechanism provides a mechanical explanation for the transient force dip observed in our hypotonic experiments. In the phase diagram (Fig. 8b (ii)), this regime maps to the deep blue zones, where higher expansion velocities induce a sharper decline in *ε*_*a*_ and drive *T*_edge_ further into the force decay region. Conversely, in the case of fast shrinkage (Case iv, *v*\* *<* 0), the negative velocity generates a kinematically strengthening effect that effectively amplifies *ε*_*a*_ (Fig. 3c), thereby triggering a counter-intuitive spike in traction force according to Eq. (39) (represented by the intensifying red gradients in Fig. 8b (iv)).

The zero-isocline 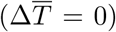 exhibits a distinct negative slope in the *ξ*-*v*\* plane, indicating that the critical velocity *v*\* shifts towards negative values as *ξ* increases. This behavior is rationalized by the traction force analysis in Eq. (36). By enforcing the equilibrium condition d*T/*d*t* = 0 and substituting the linear geometric coupling 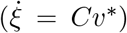 alongside the impulse acceleration approximation 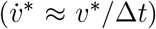, the balance relation simplifies from 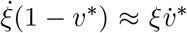 to a linear relation for the critical boundary velocity, denoted as 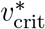:

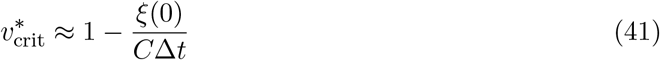

Mathematically, this equation explicitly yields a negative slope. Physically, this implies a dynamic trade-off: rapid expansion (*v*\* *>* 0) induces significant traction decay, which necessitates effective geometric strengthening (viable at small *ξ*) to maintain equilibrium. Conversely, rapid contraction (*v*\* *<* 0) induces traction accumulation, which must be counterbalanced by geometric relaxation (dominating at large *ξ*).

This disparity underscores a fundamental asymmetry between cell spreading and shrinking dynamics. Beyond this asymmetry, a pivotal insight emerges from the system’s temporal response: the Zero-Isocline exhibits a pronounced rightward shift as the observation interval Δ*t* increases. In the instantaneous limit (small Δ*t*, Fig. 8b, Fig. S6a), the stability region is confined to minimal adhesion sizes (*ξ ≪* 0.5). However, over extended intervals (Fig. S6b,c), this region expands significantly to accommodate larger adhesions (*ξ >* 3). This shift is explicitly predicted by Eq. (41), where the kinematic penalty scales inversely with Δ*t*. Physically, this stems from the distinct temporal signatures of the competing mechanisms: kinematic decay is immediate and transient, triggering traction loss upon rapid expansion, whereas geometric strengthening is cumulative and gradual (Eq. (40)). Consequently, a wider integration window allows the geometric increment Δ*ξ* to accrue sufficient magnitude to override the instantaneous kinematic decay.

The proposed kinematic-geometric competition offers a mechanistic interpretation for interpreting cellular traction under varying dynamic conditions. It is consistent with the osmotic contrast–traction loss in hypotonic vs. gain in hypertonic conditions (15)–and also recovers classical predictions for traction generation during quasi-static cell spreading (53). We therefore view the interpretation as a useful mechanical basis for studying dynamic traction responses.

### 3.3 Influence of Osmotic Shock Rate and Amplitude

To explicitly link the phase analysis with osmotic dynamics, we prescribe a specific temporal profile for the external osmotic pressure. The transition process is modeled using a standard logistic sigmoid function, *S*(*t*; *τ*), to capture the finite time required for fluid exchange and solution mixing. This profile is parameterized by the shock duration *τ* (see section S2.8 for mathematical details). Consequently, Π_out_(*t*) is defined by scaling this transition function:

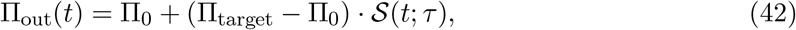

where Π_0_ is the initial pressure and Π_target_ denotes the final osmotic pressure. In this formulation, *τ* acts as an inverse proxy for the shock speed (*v*_shock_ ∝ 1*/τ*). For dimensionless analysis, we define the normalized target osmotic pressure as 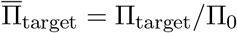.

By incorporating the Π_out_(*t*) from Eq. (42) into our simulation, we analyze the sensitivity of osmotic speed and amplitude of the dynamic traction force response. Fig. 9(a-b) quantify the peak edge velocity (*v*\*) and traction variation 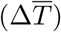 as functions of the normalized target osmotic pressure 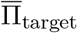. The system exhibits a monotonic dependence on shock amplitude in this model, consistent with kinematic modulation of active contractility. Specifically, hypotonic shocks 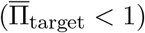 induce expansion (*v*\* *>* 0), driving a traction decay 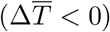 that increases with shock magnitude due to the reduction of active strain *ε*_*a*_ (kinematically dominated regime, see Fig. 8 (ii), Fig. 3d). Conversely, hypertonic shocks 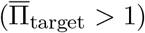 trigger contraction (*v*\* *<* 0), where larger magnitudes yield stronger kinematic amplification of *ε*_*a*_, resulting in increased traction forces (Fig. 8 (iv), Fig. 3c).

**Figure 9:**
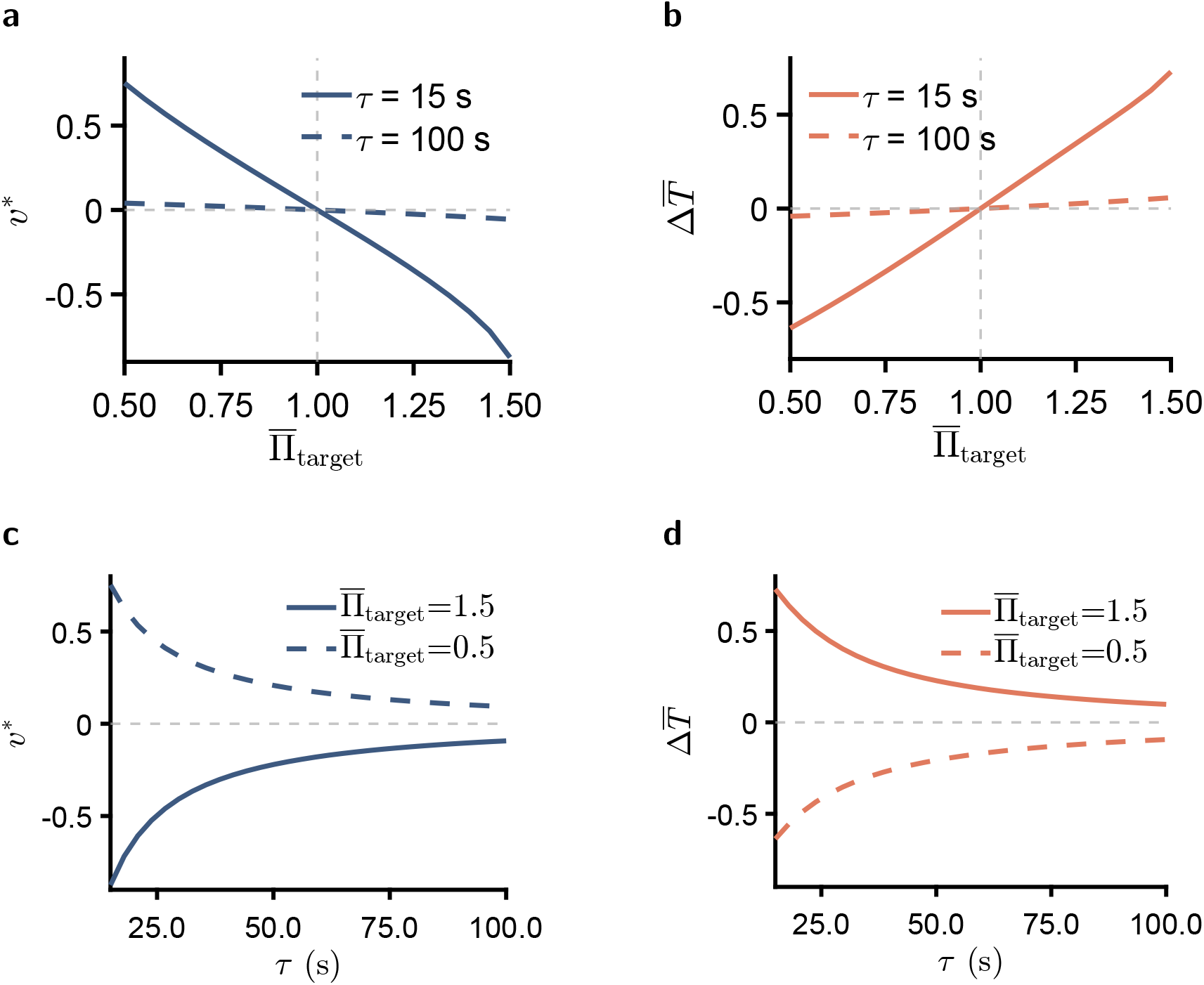
Parametric analysis of osmotic shock speed (*v*_shock_ ∝ 1*/τ*) and amplitude. (a, b) Dependence of peak edge velocity *v*\* and traction force variation 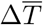 on the normalized target osmotic pressure 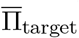. The mechanical response intensifies with the shock amplitude: stronger hypotonicity 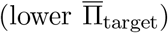 induces greater expansion and traction reduction, whereas higher hypertonicity 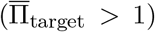 triggers enhanced contraction and traction strengthening. (c, d) Peak responses of *v*\* and 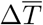 as a function of shock duration *τ* for fixed osmotic amplitudes (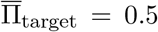 and 1.5). The magnitude of the mechanical response attenuates as the shock speed decreases (i.e., increasing *τ*).

Furthermore, in Fig. 9(a-b), for a fixed osmotic amplitude, a smaller characteristic time *τ* (i.e., a faster shock) amplifies the magnitude of both *v*\* and Δ*T*. This indicates that increasing the osmotic loading rate induces larger variations in *ε*_*a*_, thereby triggering a more pronounced transient response in traction force.

To generalize this finding beyond discrete cases, we performed a continuous parametric scan of the shock duration *τ* ranging from 15 s to 100 s. Fig. 9(c-d) map the peak responses of *v*\* and 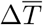 for fixed osmotic shock amplitudes (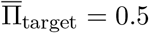 and 1.5) as a function of *τ*. As illustrated in Fig. 9c, *v*\* exhibits a hyperbolic-like decay with increasing *τ* (corresponding to decreasing osmotic shock speed). Consequently, the traction force in Fig. 9d mirrors this kinematic trend. In the hypotonic case (dashed line), a short *τ* (*<* 50 s) triggers a pronounced traction loss; however, as *τ* increases, this loss rapidly attenuates, stabilizing near zero for *τ* ≈ 100 s. Similarly, in the hypertonic case (solid line), the traction gain is concentrated at small *τ*, showing a surge that decays as the shock becomes more gradual.

Physically, higher rates and amplitudes of osmotic shock lead to faster temporal variations in cell size, represented here by *ξ*. This rapid geometric evolution generates larger excursions in the normalized edge velocity *v*\*, which in turn modulates the active strain *ε*_*a*_ and shapes the traction response in the kinematically dominated regime. To further examine whether this rate-dependent trend is robust to model-parameter variations, we conducted additional sensitivity and uncertainty analyses by varying the retrograde-flow velocity parameter *v*_*p*_, the active-strain parameter, the shock duration *τ*, and the osmotic-shock amplitude 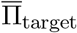; these results are summarized in Supplementary Section section S3.1. Together, these calculations indicate that rapid or large-amplitude osmotic perturbations consistently produce more pronounced changes in traction-force dynamics.

## 4 Conclusion

In summary, this study developed a dynamic biophysical framework for cellular mechano-adaptation. By treating the cell as a Hill-type active material, our theory provides an interpretation of the counter-intuitive observation that traction forces can decay during rapid expansion, and it clarifies how the relationship between cell size and force depends on dynamic loading conditions. Beyond the traditional view that mechanotransduction is primarily shaped by static adhesion geometry, our analysis highlights a geometric–kinematic competition, in which adhesion growth and edge-motion-induced active-strain suppression jointly regulate traction evolution in a perturbation-rate-dependent manner. Furthermore, the model analysis suggests that cortical stiffness contributes to osmotic recovery kinetics, with stiffer cells supporting faster geometric recoil and hydrostatic recovery. This stiffness-associated divergence may be useful for interpreting cell-type-dependent osmotic responses, alongside other regulatory mechanisms.

To calculate traction evolution, we summarized a phase diagram containing four dynamic regimes, showing whether forces amplify or decay as a function of the adhesion geometry and the kinematic rate. The phase diagram captures the osmotic paradox, namely traction loss under hypotonic expansion versus traction gain under hypertonic shrinkage, while recovering the classical prediction of traction generation during quasi-static cell spreading. The model further suggests that cell adhesion and traction responses are sensitive to both the rate and amplitude of osmotic perturbations, leading to distinct biophysical outcomes under different environmental conditions.

### Outlook

Biophysical extensions could be further integrated with biological regulation during osmotic adaptation. Processes such as cortex remodeling, adhesion turnover, ion-channel-dependent transport, and regulatory volume decrease may modulate both traction dynamics and longer-time recovery. Direct measurements of cortical organization, adhesion remodeling, and transport activity would further clarify how hypotonic swelling transiently weakens traction generation and why traction may fail to fully recover during the later response, as mechanical recoil and biochemical regulation jointly reshape osmotic cell mechanics.

## Author Contributions

**Jiarui Gan:** Conceptualization, Methodology, Software, Writing - original draft. **Xiapeng Wang:** Investigation. **Wenjie Wu:** Data curation, Validation. **Qingchuan Zhang:** Formal analysis. **Shubo Zhang:** Writing - review & editing. **Shangquan Wu:** Writing - review & editing.

## Acknowledgments

This work was financially supported by the National Natural Science Foundation of China (Grant Nos. 12232017, 12222212).

## Data Availability

The computational implementation, parameter files, sensitivity and uncertainty analysis files, processed simulation outputs, and figure-generation workflow are available at https://github.com/baartscronquist-crypto/Single-Adherent-Cell-Mechanics-in-Osmotic-Shock. Processed experimental data used for the model-experiment comparisons are included with the revision package or are deposited with the public code repository before publication. Raw microscopy and traction-force datasets are available from the corresponding authors upon reasonable request.

