## Supplementary Materials for "Kinematically-Dominated Regime Shapes Cell Traction Force Dynamics under Osmotic Shock"

### Supplemental Materials for: Kinematically-Dominated Regime Shapes Cell Traction Force Dynamics under Osmotic Shock

#### S1 Methods

##### S1.1 Cell Culture and Preparation

Five cell lines were used in the experiments. MCF-10A was maintained in their respective specific media (Procell), while MDA-MB-231 was cultured in standard RPMI-1640 or DMEM media supplemented with 10% FBS and 1% penicillin-streptomycin (Gibco). All cultures were kept at 37°C in a 5% $CO_2$ incubator. To replicate the initial conditions of the theoretical model, cells were detached using 0.25% trypsin-EDTA and seeded onto hydrogel substrates. They were allowed to settle and establish weak adhesion for 20 minutes prior to the application of osmotic shocks.

##### S1.2 Cortical Stiffness Measurement (AFM)

The Young's modulus of the cell cortex was characterized using Atomic Force Microscopy (Nanowizard Sense+, Bruker) mounted on an inverted microscope. We employed silicon nitride cantilevers (DNP-D, nominal $k = 0.06$ N/m) equipped with a pyramidal tip (20 nm radius). The spring constants were calibrated via the Sader method. To specifically probe the cortical stiffness rather than the deep cytoplasm, indentation experiments were performed in contact mode with low setpoints (0.4–0.6 nN), ensuring indentation depths remained below 1 $\mu$m. The resulting force-indentation curves were fitted to the Hertz-Sneddon model to extract the elastic modulus.

##### S1.3 Traction Force Microscopy (TFM)

Traction forces were quantified on 10kPa hydrogels embedded with fluorescent marker beads. Briefly, hydrogel surfaces were activated using EDC/NHS chemistry to promote bead attachment (0.5 $\mu$m, carboxyl-modified) and subsequent cell adhesion. The activation mixture contained 0.5% (w/v) NHS and 2% (w/v) EDC. Following washing steps, cells were seeded and cultured for 20 minutes before imaging. We acquired Z-stack images of the fluorescent beads using confocal microscopy. Deformation fields were determined by comparing bead positions in the stressed state (with cells) and the relaxed state (after 0.5% SDS-induced cell detachment) using Digital Image Correlation (PMLAB). The displacement field was then inverted to compute the traction stress field using the open-source PYTFM package, which solves the inverse Boussinesq problem in Fourier space. The mean traction force magnitude was calculated for statistical analysis.

##### S1.4 Model Geometry Schematic

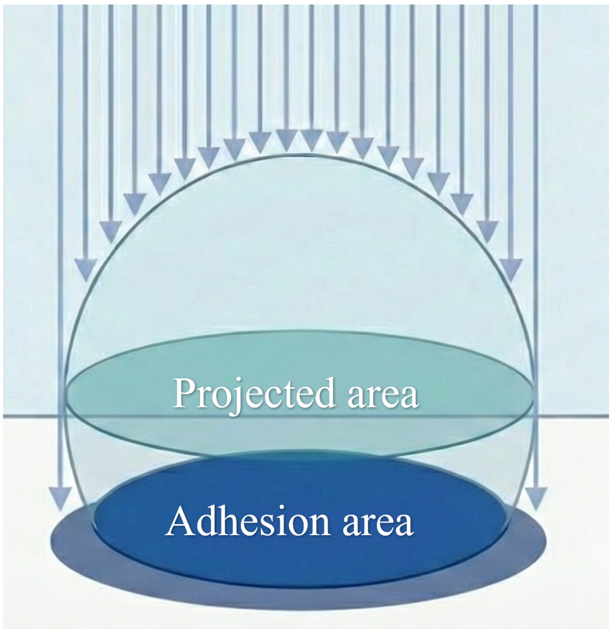


Figure S1: Schematic illustrating the difference between projected area and adhesion area.

### S2 Analytical and numerical Derivations

#### S2.1 Derivation of the Initial Mechano-chemical Equilibrium Constraint

To determine the physically admissible initial state of the cell, we enforce the steady-state conditions for both water and ion fluxes, coupled with the mechanical equilibrium of the membrane.

##### S2.1.1 Reduction to Scalar Formulation.

As described in the main text, the cortex is modeled as an elastic thin shell described by a tensorial constitutive law. To facilitate the analytical derivation of the initial equilibrium, we simplify the cell geometry to an axisymmetric model subject to isotropic inflation. Consequently, the surface stress state is assumed to be equibiaxial and planar. Under this assumption, the stress and strain tensors reduce to scalar forms: $\boldsymbol{\sigma}_{\text{cort}} = \sigma \mathbf{I}$ and $\boldsymbol{\varepsilon}_{\text{cort}} = \varepsilon \mathbf{I}$, where $\mathbf{I}$ is the 2D identity tensor.

Substituting these scalar forms into constitutive laws, the trace term becomes $\text{tr}(\boldsymbol{\varepsilon}_{\text{cort}}) = 2\varepsilon$. The scalar constitutive relation thus simplifies to:

$$\sigma = \frac{E_m}{1 - \nu_m} \varepsilon + \sigma_{\text{act}}, \tag{S1}$$

where $\sigma$ represents the isotropic membrane tension and $\sigma_{\text{act}}$ is the magnitude of the active stress. This scalar formulation is utilized in the following derivation.

##### S2.1.2 Water Flux Equilibrium.

The system is initially in a steady state where the net water flux $J_w$ vanishes. According to the Starling equation, the water flux is driven by the imbalance between the osmotic pressure difference $\Delta\Pi$ and the hydrostatic pressure difference $\Delta P$:

$$J_w = \alpha(\Delta\Pi - \Delta P) = 0 \implies \Delta P = \Delta\Pi, \tag{S2}$$

where $\alpha$ is the hydraulic permeability.

##### S2.1.3 Ion Flux Equilibrium.

Simultaneously, the net ion flux $dn/dt$ must be zero. The total ion flux comprises active ($J_a$) and passive ($J_p$) components. Introducing the mechano-sensitive gating formulations, the equilibrium implies a balance between the osmotic potential and the membrane tension $\sigma$:

$$\gamma(\Delta\Pi_c - \Delta\Pi) - \beta(\sigma - \sigma_c)\Delta\Pi = 0, \tag{S3}$$

where $\gamma$ and $\beta$ are the chemical and mechanical gating coefficients, respectively, and definitions of critical thresholds $(\Delta\Pi_c, \sigma_c)$ follow the standard mechanobiological models.

##### S2.1.4 Mechanical Equilibrium (Laplace's Law).

For the thin-walled structure, the hydrostatic pressure difference $\Delta P$ is balanced by the scalar membrane tension $\sigma$ according to the generalized Laplace's law:

$$\Delta P = \frac{2\sigma d}{R}, \tag{S4}$$

where $d$ is the effective thickness of the cortex and $R$ is the radius of curvature.

##### S2.1.5 Geometric Coupling Constraint.

By combining Eq. (S2) ($\Delta P = \Delta\Pi$) with Eq. (S3) and Eq. (S4), we eliminate the pressure terms to derive an explicit relationship between the radius $R$ and the tension $\sigma$:

$$R(\sigma) = \frac{2\sigma d}{\gamma\Delta\Pi_c}\left[\beta(\sigma - \sigma_c) - \gamma\right]. \tag{S5}$$

Finally, substituting the geometric definition of strain $\varepsilon(R, h)$ into the scalar constitutive law (Eq. (S1)) yields $\sigma$ as a function of geometry $\sigma(R, h)$. Inserting this back into Eq. (S5) leads to the implicit constraint:

$$f(R, h) = R - R\big(\sigma(R, h)\big) = 0. \tag{S6}$$

This confirms that the initial geometric parameters $\{R, h\}$ are coupled and must lie on the solution manifold of Eq. (S6).

#### S2.2 Derivation of the power balance

The total energy of the system is denoted as $\mathcal{U}$, which is decomposed into the cortical strain energy ($\mathcal{U}_{\text{cort}}$) and the residual contributions ($\mathcal{U}_{\text{res}}$) comprising adhesion chemical, adhesion elastic and stress fiber energies:

$$\mathcal{U} = \mathcal{U}_{\text{cort}} + \mathcal{U}_{\text{res}}. \tag{S7}$$

In the main text, the power balance is driven by the hydrostatic pressure difference $\Delta P$. Physically, this pressure represents the mechanical restoring force generated by the cell cortex. Therefore, we identify $\Delta P$ as the mechanical pressure ($P_{\text{mech}}$), defined as the work-conjugate variable to the cortical deformation:

$$\Delta P \equiv P_{\text{mech}} = \frac{\partial \mathcal{U}_{\text{cort}}}{\partial V}. \tag{S8}$$

Substituting this definition into the global power balance equation (In Main Text, $\frac{\mathrm{d}\mathcal{U}}{\mathrm{d}t} = \Delta P \frac{\mathrm{d}V}{\mathrm{d}t}$), we obtain:

$$\frac{\mathrm{d}}{\mathrm{d}t}(\mathcal{U}_{\text{cort}} + \mathcal{U}_{\text{res}}) = \left(\frac{\partial \mathcal{U}_{\text{cort}}}{\partial V}\right)\frac{\mathrm{d}V}{\mathrm{d}t}. \tag{S9}$$

Applying the chain rule to the cortical energy term, we have $\frac{\mathrm{d}\mathcal{U}_{\text{cort}}}{\mathrm{d}t} = \frac{\partial \mathcal{U}_{\text{cort}}}{\partial V}\frac{\mathrm{d}V}{\mathrm{d}t}$. Consequently, the cortical terms on both sides of the equation cancel out, yielding:

$$\frac{\mathrm{d}\mathcal{U}_{\text{res}}}{\mathrm{d}t} = 0. \tag{S10}$$

Thus, our power balance formulation mathematically enforces that the residual energy components remain constant during the simulation steps. This is consistent with the physiological fact that focal adhesion remodeling operates on a significantly longer timescale compared to the rapid osmotic response.

#### S2.3 Derivation of the Dynamic Traction Force Model

To quantify the traction force response under dynamic conditions, we establish a coupling framework linking macroscopic edge kinetics to microscopic molecular binding. We introduce the dimensionless edge velocity $v^* = v/v_p$, normalized by the characteristic polymerization velocity $v_p$. The total traction stress, $T$, is defined as the product of the closed bond density, $\rho$, and the force borne by individual bonds, $F_b$:

$$T(\xi, v^*) = \rho(\xi, v^*) \cdot F_b(\xi, v^*). \tag{S11}$$

**Elasticity and Kinematic Coupling.** We employ a shear-lag model involving Bessel functions to describe the displacement field. The edge displacement $u(r_c)$ is analytically given by:

$$u(r_c) = \varepsilon_a(v^*)(1+\nu)\lambda \cdot \mathcal{F}(\xi), \quad \text{with} \quad \mathcal{F}(\xi) = \frac{\xi I_1(\xi)}{\xi I_0(\xi) - (1-\nu) I_1(\xi)}, \tag{S12}$$

where $\xi = r_c/\lambda$ is the dimensionless cell size. To capture the friction of actomyosin motors, the active contractile strain $\varepsilon_a(v^*)$ decays with the normalized velocity $v^*$ according to a phenomenological relation:

$$\varepsilon_a(v^*) = \varepsilon_{\mathrm{a}}(0)(1 - v^*). \tag{S13}$$

Assuming linear spring behavior for adhesion bonds with stiffness $k_{\text{bond}}$, the single bond force is expressed as $F_b(\xi, v^*) = k_{\text{bond}}|u(r_c)|$.

**Kinetics and Bond Density.** The bond density evolves according to the reversible binding kinetics between free receptors (Rec) and ligands (Lig):

$$\text{Rec} + \text{Lig} \underset{k_{\text{off}}}{\overset{k_{\text{on}}}{\rightleftharpoons}} \text{Bond}. \tag{S14}$$

Let $\rho_R^{\text{tot}}$ and $\rho_L^{\text{tot}}$ denote the total surface densities of receptors and ligands, respectively. Mass conservation implies that the densities of available free species are $(\rho_R^{\text{tot}} - \rho)$ and $(\rho_L^{\text{tot}} - \rho)$. Consequently, the reaction rate equation is given by:

$$\frac{\mathrm{d}\rho}{\mathrm{d}t} = k_{\text{on}}(\rho_R^{\text{tot}} - \rho)(\rho_L^{\text{tot}} - \rho) - k_{\text{off}}(F_b)\rho. \tag{S15}$$

Assuming that chemical equilibrium is reached instantaneously relative to the dynamics of cell-substrate mechanical interactions, we set $\frac{\mathrm{d}\rho}{\mathrm{d}t} = 0$. This yields a quadratic equation for the steady-state bond density:

$$\rho^2 - (\Sigma_{\text{tot}} + K_D)\rho + \rho_R^{\text{tot}}\rho_L^{\text{tot}} = 0, \tag{S16}$$

where $\Sigma_{\text{tot}} = \rho_R^{\text{tot}} + \rho_L^{\text{tot}}$, and $K_D(F_b) = k_{\text{off}}(F_b)/k_{\text{on}}$ is the force-dependent dissociation constant determined by the catch-slip model. To ensure physical validity ($\rho \leq \min(\rho_R^{\text{tot}}, \rho_L^{\text{tot}})$), we select the smaller root:

$$\rho(r) = \frac{(\Sigma_{\text{tot}} + K_D) - \sqrt{(\Sigma_{\text{tot}} + K_D)^2 - 4\rho_R^{\text{tot}}\rho_L^{\text{tot}}}}{2}. \tag{S17}$$

#### S2.4 Asymptotic Analysis of Bond Density $\rho(\xi)$

In this section, we derive the analytical limits of the bond density $\rho(\xi)$ in the small adhesion regime ($\xi \to 0$) and the large adhesion saturation regime ($\xi \to \infty$).

**Regime I: small adhesion ($\xi \to 0$)**

For small $\xi$, we assume a linear kinematic approximation $u(\xi) \approx \varepsilon_a \lambda \xi$. Consequently, the single bond force follows $F_b(\xi) \approx k_{\text{sub}} \varepsilon_a \lambda \xi$. Linearizing the dissociation constant $K_D$ for small forces yields:

$$K_D(\xi) \approx K_D^0 - \left( \frac{\chi k_{\text{sub}} \varepsilon_a \lambda}{k_{\text{on}}} \right) \xi, \tag{S18}$$

where $\chi$ is the effective mechanosensitivity coefficient derived from the catch-slip parameters. By expanding Eq. (S17) around the equilibrium density $\rho_0 = \rho(K_D^0)$, we obtain the asymptotic profile:

$$\rho(\xi) \approx \rho_0 - C_0 \varepsilon_a \lambda \xi, \quad \text{with} \quad C_0 = \mathcal{S}_\rho \frac{\chi k_{\text{sub}}}{k_{\text{on}}}, \tag{S19}$$

where $\mathcal{S}_\rho$ is a sensitivity factor. This indicates that the bond density initially decreases linearly with $\xi$ due to force-induced unbinding.

**Regime II: large adhesion saturation ($\xi \to \infty$)**

In the limit of large $\xi$, the displacement approaches a saturation value $u_\infty \approx \varepsilon_a (1 + \nu) \lambda$. The bond tension caps at $F_{\text{sat}} = k_{\text{sub}} u_\infty$, leading to a constant dissociation rate $K_D^\infty$. Consequently, the bond density converges to a stable plateau $\rho_\infty$ rather than vanishing. This reflects a steady-state adhesion limited by the maximum contractility of the cytoskeleton.

#### S2.5 Traction Force Monotonicity Analysis

To analyze the dependence of the traction force evolution rate on the dimensionless velocity $v^*$, we start with the approximation derived in the main text:

$$\frac{\mathrm{d}T_{\text{edge}}}{\mathrm{d}t} \approx C_1 \dot{\xi} (1 - v^*), \tag{S20}$$

where $C_1$ is a positive material constant. The time-dependent normalized radius $\xi(t)$ is defined by the integral relation:

$$\xi(t) = \xi_0 + \frac{v_{\text{p}}}{\lambda} \int_0^t v^*(t') \, \mathrm{d}t'. \tag{S21}$$

Differentiating $\xi(t)$ with respect to time yields the relationship between the geometric expansion rate and the instantaneous velocity:

$$\dot{\xi} = \frac{\mathrm{d}\xi}{\mathrm{d}t} = \frac{v_{\text{p}}}{\lambda} v^*. \tag{S22}$$

Substituting Eq. (S22) into Eq. (S20), we express the traction rate $\mathcal{R}$ solely as a function of $v^*$:

$$\mathcal{R}(v^*) \equiv \frac{\mathrm{d}T_{\text{edge}}}{\mathrm{d}t} \approx \left( \frac{C_1 v_{\text{p}}}{\lambda} \right) v^* (1 - v^*) = K v^* (1 - v^*), \tag{S23}$$

where $K = C_1 v_{\text{p}} / \lambda > 0$ is a strictly positive coefficient. To determine the monotonicity of $\mathcal{R}$ with respect to $v^*$, we examine its first derivative:

$$\frac{\partial \mathcal{R}}{\partial v^*} = K \frac{\mathrm{d}}{\mathrm{d}v^*} (v^* - v^{*2}) = K(1 - 2v^*). \tag{S24}$$

The sign of the derivative depends on the magnitude of $v^*$, leading to two distinct regimes:

1. **Monotonically Increasing Regime ($v^* < 0.5$):** When $v^* < 0.5$, $\frac{\partial \mathcal{R}}{\partial v^*} > 0$. In this low-velocity regime, the geometric expansion term ($\dot{\xi}$) dominates. Consequently, an increase in velocity leads to a higher rate of traction accumulation ($\mathrm{d}T_{\text{edge}}/\mathrm{d}t$ increases).

2. **Monotonically Decreasing Regime ($v^* > 0.5$):** When $v^* > 0.5$, $\frac{\partial \mathcal{R}}{\partial v^*} < 0$. In this high-velocity regime, the frictional slip factor $(1 - v^*)$ dominates. As $v^*$ increases further towards unity, the diminishing friction efficiency suppresses the force generation, causing $\mathrm{d}T_{\mathrm{edge}}/\mathrm{d}t$ to decrease despite the faster expansion.

Thus, the traction rate is not globally monotonic; it exhibits a parabolic profile with a peak at $v^* = 0.5$. The positive correlation between velocity and traction growth rate holds strictly only when $v^* \in [0, 0.5)$.

#### S2.6 Dynamic DAE Solver Strategy: Dimensionality Reduction via Geometric Substitution

To efficiently solve the coupled geometric and hydraulic evolution from time step $n$ to $n+1$, we employ a dimensionality reduction strategy. Instead of simultaneously solving for the independent variables radius $R_{n+1}$ and height $h_{n+1}$, we introduce an auxiliary variable $\eta$ representing the total geometric span:

$$\eta = R_{n+1} + h_{n+1}. \tag{S25}$$

Under the ansatz of a fixed $\eta$ in the outer solver loop, the volume-conservation equation can be decoupled and reduced to an analytical quadratic solution for $R_{n+1}$.

##### S2.6.1 Geometric Linearization

The primary advantage of the transformation $h_{n+1} = \eta - R_{n+1}$ is that it linearizes the highly nonlinear geometric expressions for spherical caps.

**Volume $V_{n+1}$** Substituting $h_{n+1}$ into the volume formula $V = \frac{\pi}{3}(R+h)^2(2R-h)$ yields:

$$\begin{aligned} V_{n+1}(\eta, R_{n+1}) &= \frac{\pi}{3}\eta^2 \left[2R_{n+1} - (\eta - R_{n+1})\right] \\ &= \frac{\pi}{3}\eta^2 (3R_{n+1} - \eta) \\ &= \underbrace{(\pi\eta^2)}_{K_V} R_{n+1} - \underbrace{\frac{\pi}{3}\eta^3}_{C_V}. \end{aligned} \tag{S26}$$

Crucially, for a fixed $\eta$, the volume $V_{n+1}$ becomes a linear function of $R_{n+1}$.

**Surface Area $A_{n+1}$** Similarly, the surface area $A = 2\pi R(R+h)$ simplifies to:

$$A_{n+1}(\eta, R_{n+1}) = 2\pi R_{n+1}(\eta) = \underbrace{(2\pi\eta)}_{K_A} R_{n+1}. \tag{S27}$$

Thus, the surface area also scales linearly with $R_{n+1}$ for a constant $\eta$.

##### S2.6.2 Quadratic Form of Hydraulic Flux

The volume conservation requires that the change in volume equals the integrated flux:

$$V_{n+1} - V_n = \Delta V_{\mathrm{flux}}. \tag{S28}$$

The flux term is defined by $\Delta V_{\mathrm{flux}} \approx \Delta t \cdot J_v \cdot A_{n+1}$. The water flux density $J_v$, described by the Kedem-Katchalsky equation, depends on the Laplace pressure ($\Delta P \propto 1/R$). By performing a local linearization of the pressure term around the previous state $R_n$, the flux density can be approximated as a linear function of $R_{n+1}$:

$$J_v(R_{n+1}) \approx \alpha R_{n+1} + \beta, \tag{S29}$$

where $\alpha$ and $\beta$ are coefficients derived from the state at step $n$.

Multiplying this linearized flux density by the linearized area (Eq. S27), the total volume flux becomes **quadratic** in $R_{n+1}$:

$$\begin{aligned}\Delta V_{\text{flux}} &= \Delta t \cdot (\alpha R_{n+1} + \beta) \cdot (K_A R_{n+1}) \\ &= \Delta t \left[ (K_A \alpha) R_{n+1}^2 + (K_A \beta) R_{n+1} \right]. \end{aligned} \tag{S30}$$

##### S2.6.3 Analytical Resolution

Substituting the linearized volume (Eq. S26) and quadratic flux (Eq. S30) back into the conservation equation yields:

$$(K_V R_{n+1} - C_V) - V_n = \Delta t \left[ (K_A \alpha) R_{n+1}^2 + (K_A \beta) R_{n+1} \right]. \tag{S31}$$

Rearranging terms results in a standard quadratic equation for $R_{n+1}$:

$$\mathcal{A} R_{n+1}^2 + \mathcal{B} R_{n+1} + \mathcal{C} = 0, \tag{S32}$$

with coefficients determined entirely by the fixed ansatz $\eta$ and previous state values:

$$\mathcal{A} = \Delta t \cdot K_A \alpha \tag{S33a}$$

$$\mathcal{B} = \Delta t \cdot K_A \beta - K_V \tag{S33b}$$

$$\mathcal{C} = C_V + V_n \tag{S33c}$$

Solving this quadratic equation yields the explicit physical solution for the radius:

$$R_{n+1} = \frac{-\mathcal{B} + \sqrt{\mathcal{B}^2 - 4\mathcal{A}\mathcal{C}}}{2\mathcal{A}}. \tag{S34}$$

This analytical expression formally defines the function $R_{n+1} = \mathcal{F}(\eta)$, which serves as the fundamental geometric constraint for the outer solver loop.

##### S2.6.4 Solver Closure: Energy Consistency Loop

The algebraic resolution derived above establishes a deterministic mapping $R_{n+1} = \mathcal{F}(\eta)$, ensuring that volume conservation is strictly satisfied for any candidate $\eta$. Consequently, the solver task reduces from a 2D search to a 1D root-finding problem for the scalar $\eta_{n+1}$. The closure of the system is achieved by enforcing the thermodynamic power balance.

For a candidate $\eta$, once the corresponding physical radius $R(\eta)$ and height $h(\eta) = \eta - R(\eta)$ are determined, the contact radius $r_c$ is calculated as:

$$r_c(\eta) = \sqrt{R(\eta)^2 - h(\eta)^2}. \tag{S35}$$

The velocity of the contact line, $v_{\text{edge}}$, is then computed via a backward difference scheme relative to the previous step $n$:

$$v_{\text{edge}} = \frac{r_c(\eta) - r_{c,n}}{\Delta t}. \tag{S36}$$

This velocity determines the dimensionless kinetic state $v^* = v_{\text{edge}}/v_{\text{p}}$. Substituting $v^*$ into the microscopic traction model (Eq. (S11)), we obtain the velocity-dependent traction/energy term $T(\xi, v^*)$, which sets the non-cortical energy dissipation or storage term in the model.

The correct value of $\eta_{n+1}$ is identified by satisfying the global power balance equation:

$$\mathcal{R}(\eta) = \frac{\mathcal{U}_{\text{total}}(\eta) - \mathcal{U}_n}{\Delta t} - \Delta P(\eta) \frac{V(\eta) - V_n}{\Delta t} = 0. \tag{S37}$$

Here, the total energy rate $\dot{\mathcal{U}}$ explicitly incorporates the contributions from $T(\xi, v^*)$. The DAE solver iterates on $\eta$ until the residual $\mathcal{R}(\eta)$ converges to zero. Once the valid $\eta_{n+1}$ is found, the final geometric state $R_{n+1}$ is strictly defined by the quadratic mapping $\mathcal{F}(\eta_{n+1})$.

##### S2.7 Computational Algorithm Flowchart

The numerical strategy derived above is implemented according to the following workflow. The nested loop highlights the 1D semi-analytical solver.

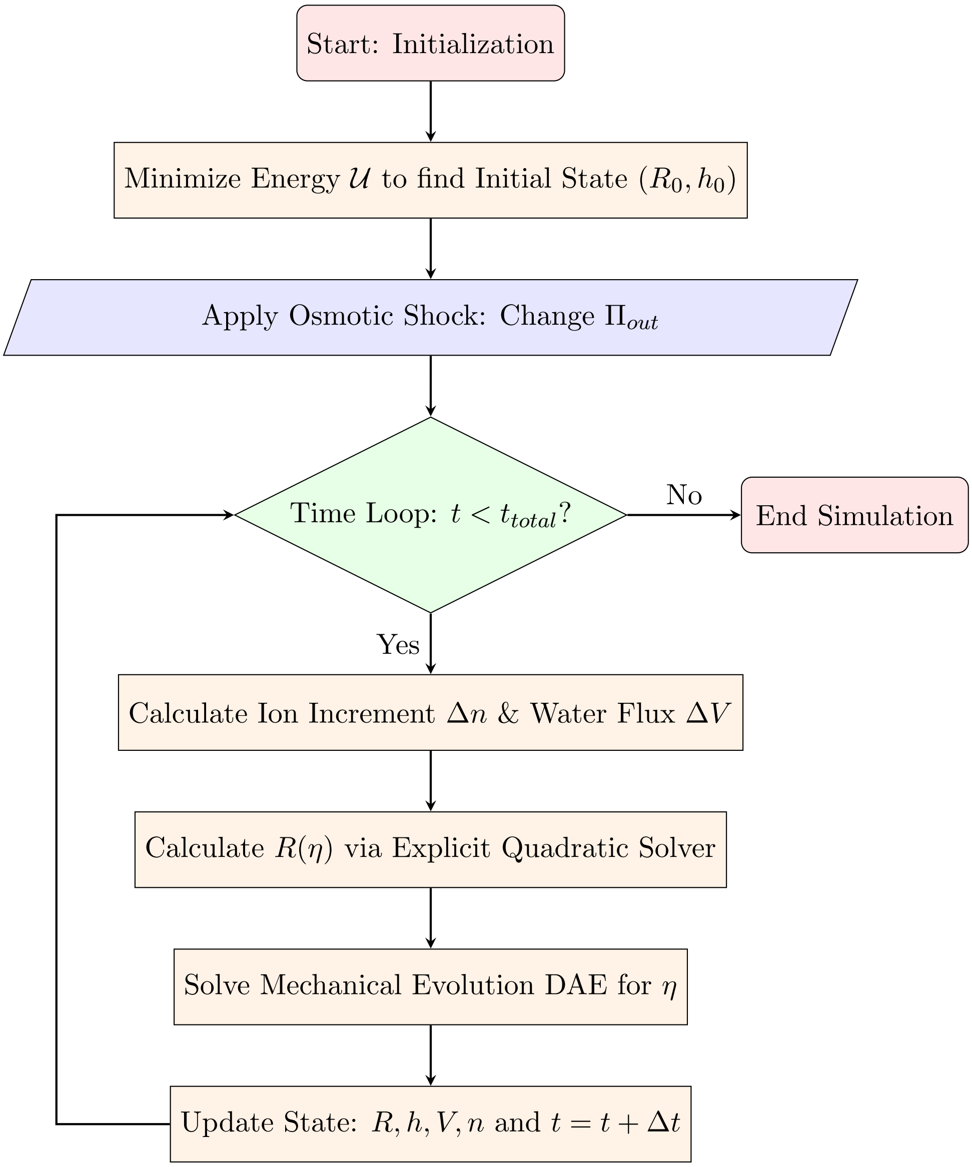


Figure S2: Flowchart of the simulation process. The diagram illustrates the initialization phase, the time-stepping loop driven by osmotic shocks, and the nested 1D semi-analytical solver strategy.

##### S2.8 Definition of logistic sigmoid function

To give different Osmotic Shock Rate and Amplitude, we prescribe the temporal profile of the external pressure. To ensure the transition is strictly controlled by the shock duration $\tau$, we define the time center $t_c$ and the steepness coefficient $k_{\text{speed}}$ as:

$$t_c = \frac{\tau}{2}, \quad k_{\text{speed}} = \frac{12}{\tau} \tag{S38}$$

Here, $t_c$ centers the shock within the interval, while the factor 12 in $k_{\text{speed}}$ is chosen to ensure that the sigmoid function effectively completes its transition (from 0.2% to 99.8%) within the range $t \in [0, \tau]$.Substituting these parameters into the standard logistic equation $\mathcal{S}(t) = [1 + \exp(-k_{\text{speed}}(t - t_c))]^{-1}$, we obtain the specific form of the normalized transition function:

$$\mathcal{S}(t; \tau) = \frac{1}{1 + \exp\left[-\frac{12}{\tau}\left(t - \frac{\tau}{2}\right)\right]} \tag{S39}$$

#### S3 Supplementary Results

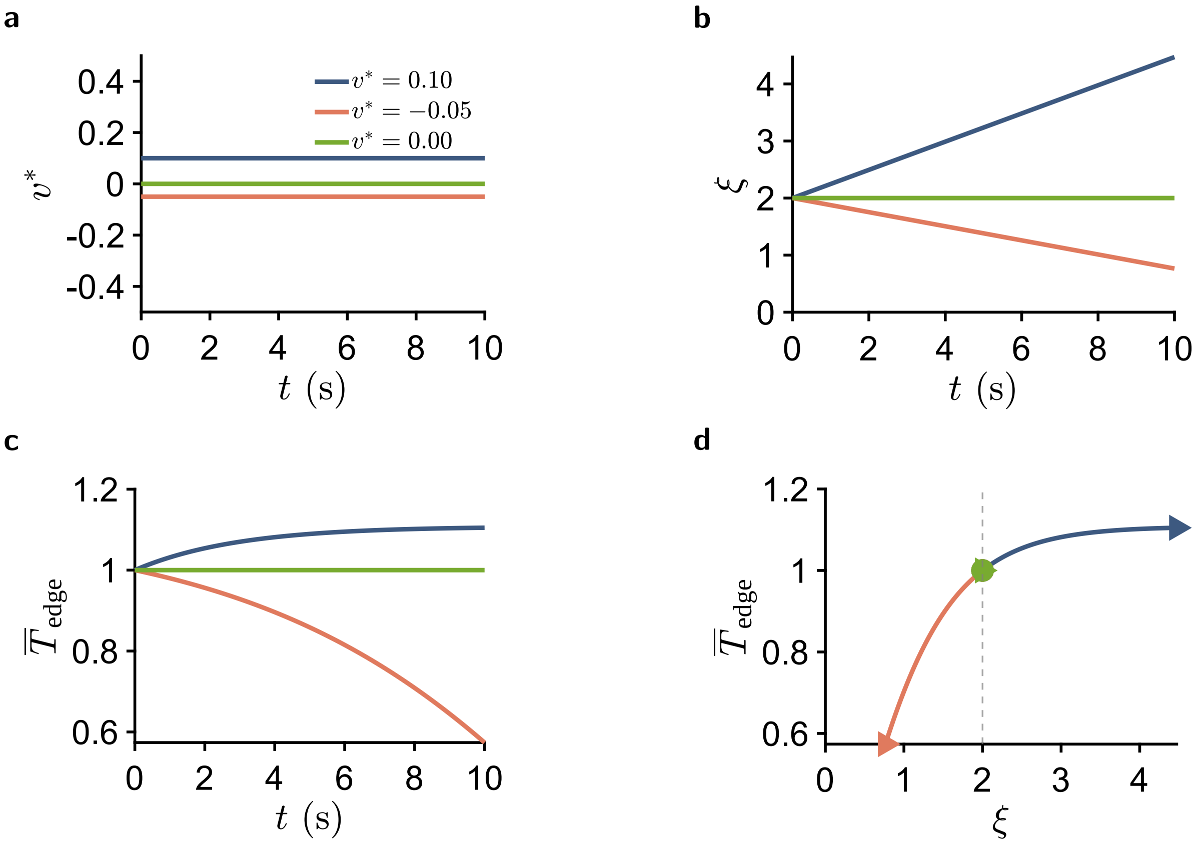


Figure S3: Representative results for the zero acceleration case ($\dot{v} = 0$). (a) Prescribed velocity profiles. (b) Corresponding time evolution of cell size $\xi$. (c) Normalized phase trajectories in the $\xi - \overline{T}$ plane. (d) Temporal dynamics of traction force, highlighting the distinct responses to expansion and contraction.

##### S3.1 Sensitivity and Uncertainty Analysis

The scanned parameters were $v_{\text{p}}$, $\varepsilon_a(0)$, $\tau$ (shock duration), and $\overline{\Pi}_{\text{target}}$ (osmotic amplitude). The perturbation ranges are listed in table S1. For computational efficiency, we set the analysis window at $t = 5\,\text{s}$.

As shown in Fig. S7, sensitivity ranking identifies $\tau$ as the most influential parameter for both the traction drop fraction (relative span $\approx 2.5$) and $\min(\overline{T})$ (relative span $\approx 0.85$). The edge-velocity scale $v_{\text{p}}$ ranks second in both metrics, followed by $\varepsilon_a(0)$ and $\overline{\Pi}_{\text{target}}$. The dominance of $\tau$ indicates that the shock time scale, rather than the osmotic amplitude alone, is the primary driver of the transient traction response.

The uncertainty analysis retained $n = 10$ parameter sets satisfying the hypotonic traction-loss branch (perturbation ranges in table S1). The trajectory in Fig. S8a yields a median traction drop of 35.3%, with $\min(\overline{T})$ ranging from 0.259 (5th percentile) to 0.848 (95th percentile), and a median final normalized traction of 0.647 (5th–95th: 0.259–0.848). We note a transient traction

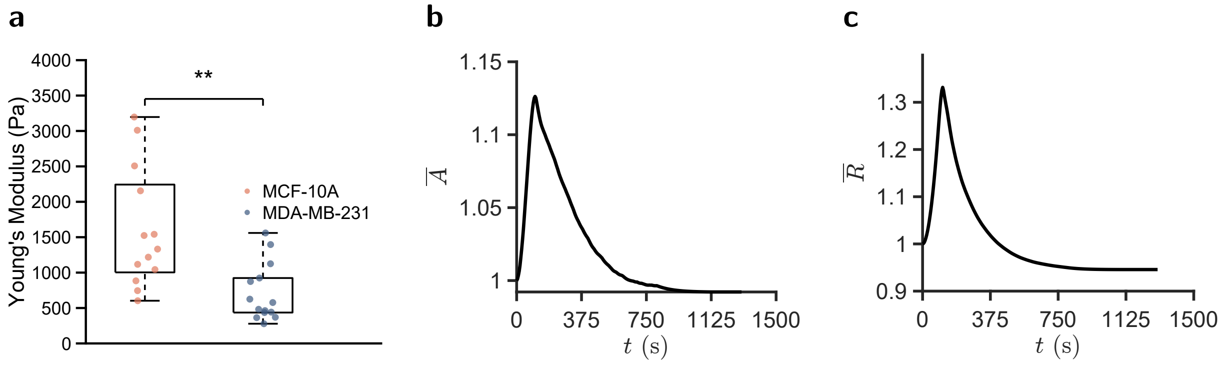


Figure S4: Experimental characterization and geometric evolution under osmotic shock in simulation. (a) Young's modulus of MCF-10A (stiff) and MDA-MB-231 (soft) cells measured by AFM ($^{**}p < 0.01$). (b) Temporal evolution of the normalized effective area ($\bar{A}$). (c) Temporal evolution of the normalized radius ($\bar{R}$). The simulation results in (b) and (c) correspond to the parameters of the stiff cell.

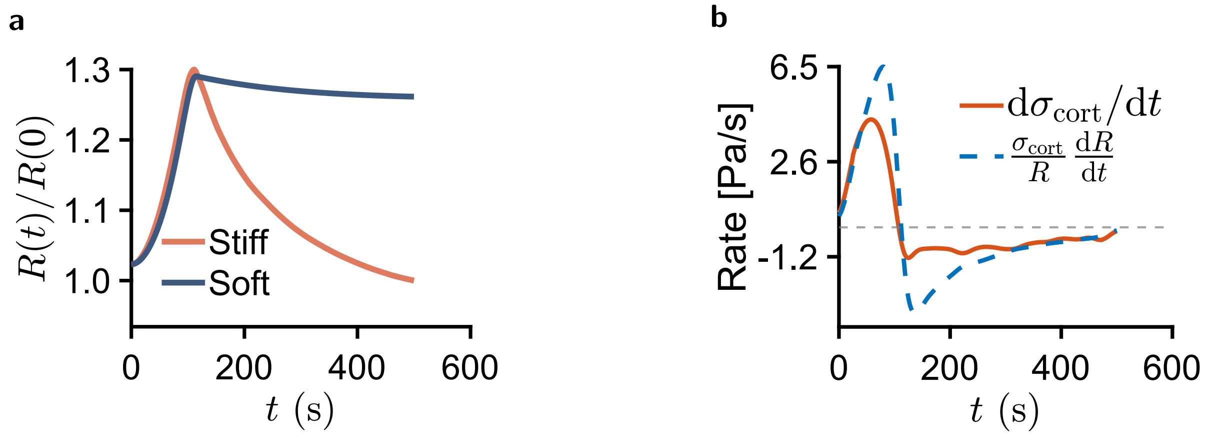


Figure S5: Temporal evolution of cell mechanics during osmotic shock. (a) Time course of the cell radius. (b) Comparison between the geometric rebound term (blue line) and the stress rebound term (red line) for a stiff cell.

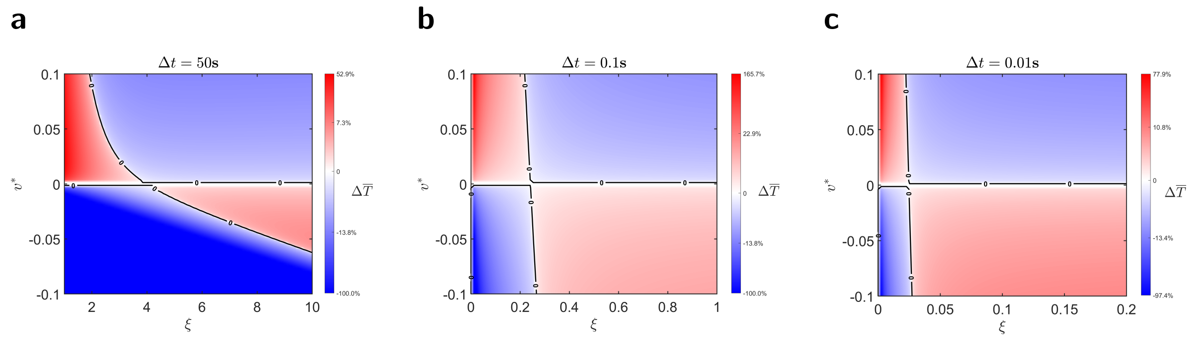


Figure S6: Effect of observation window $\Delta t$ on stability phase diagrams. (a) Long-time limit ($\Delta t = 50\,\mathrm{s}$), showing strict stability boundaries. (b) Intermediate timescale ($\Delta t = 10\,\mathrm{s}$). (c) Instantaneous limit ($\Delta t = 0.01\,\mathrm{s}$), where the instability region is significantly suppressed.

recovery at the beginning of the window, which we attribute to the initially gradual osmotic gradient (Fig. S8b).

The distributions are shown in Fig. S9: (a) drop fraction, (b) minimum normalized edge traction $\min(\overline{T})$, and (c) final normalized edge traction $\overline{T}_{\text{final}}$. For the relatively mild hypotonic shock considered here, the ensemble response is characterized by a moderate decrease in traction. The median drop fraction remains below 0.5, meaning that the typical simulated response retains more than half of the initial normalized traction at its minimum.

Table S1: Hypotonic sensitivity perturbation ranges.

| Parameter | Baseline | Sensitivity | Uncertainty |
|---|---|---|---|
| $v_\text{p}$ ($\text{m}\,\text{s}^{-1}$) | $5.0 \times 10^{-7}$ | $2.5 \times 10^{-7}$, $5.0 \times 10^{-7}$, $1.0 \times 10^{-6}$ | $U(2.5 \times 10^{-7}, 1.0 \times 10^{-6})$ |
| $\varepsilon_a(0)$ | 0.05 | 0.01, 0.05, 0.09 | $U(0.01, 0.09)$ |
| $\tau$ (shock duration) | 20 s | 15, 20, 25 s | $U(15, 25)$ s |
| $\overline{\Pi}_{\text{target}}$ | 0.45 | 0.33, 0.45, 0.58 | $U(0.33, 0.58)$ |

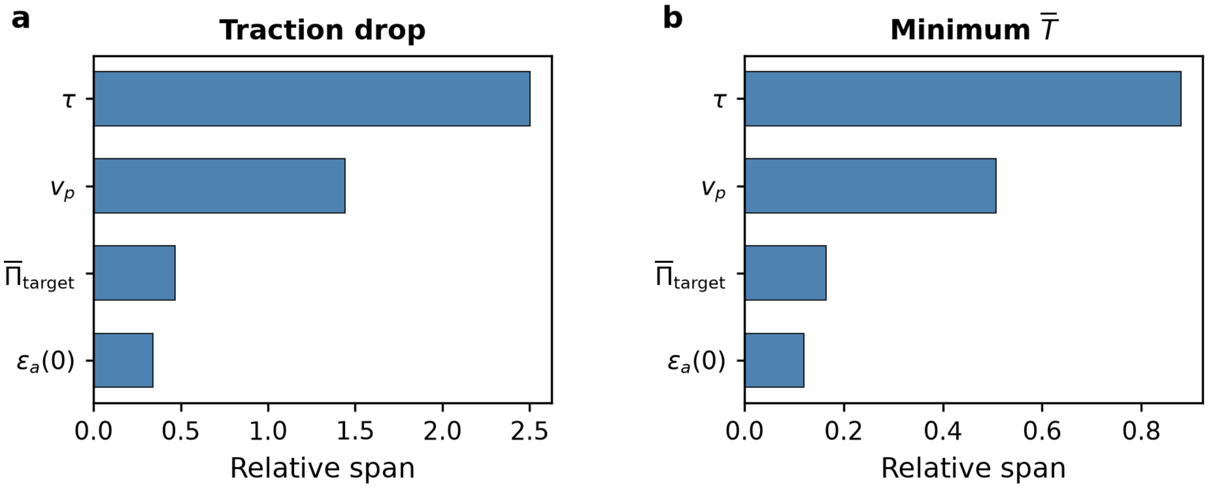


Figure S7: Sensitivity ranking of the hypotonic traction response. (a) Relative span of the traction drop fraction. (b) Relative span of the minimum normalized edge traction $\overline{T}$. Parameters are ordered by decreasing sensitivity; the most influential parameters are the shock time scale $\tau$, the osmotic amplitude $\overline{\Pi}_{\text{target}}$, the active-strain amplitude $\varepsilon_a(0)$, and the edge-velocity scale $v_\text{p}$.

#### S3.2 Poroelasticity Discussion

Poroelasticity may contribute to short-time cellular mechanical responses under osmotic perturbation, since osmotic loading involves water permeation and fluid redistribution through the cell and cortical network.

To examine a volume-related mechanical aspect relevant to this response, we added a Poisson-ratio control by varying the cortical Poisson ratio $\nu_m$ and the stress-fiber Poisson ratio $\nu_{\text{SF}}$. The results in Fig. S10 show that changing $\nu_m$ affects the normalized radius $\bar{R}$, whereas the normalized stress and normalized edge traction are only weakly affected by variations in $\nu_m$ and $\nu_{\text{SF}}$, respectively.

This behavior suggests that the volume-related mechanical response is mainly reflected in cell-size recovery and cortical deformation during osmotic water exchange.

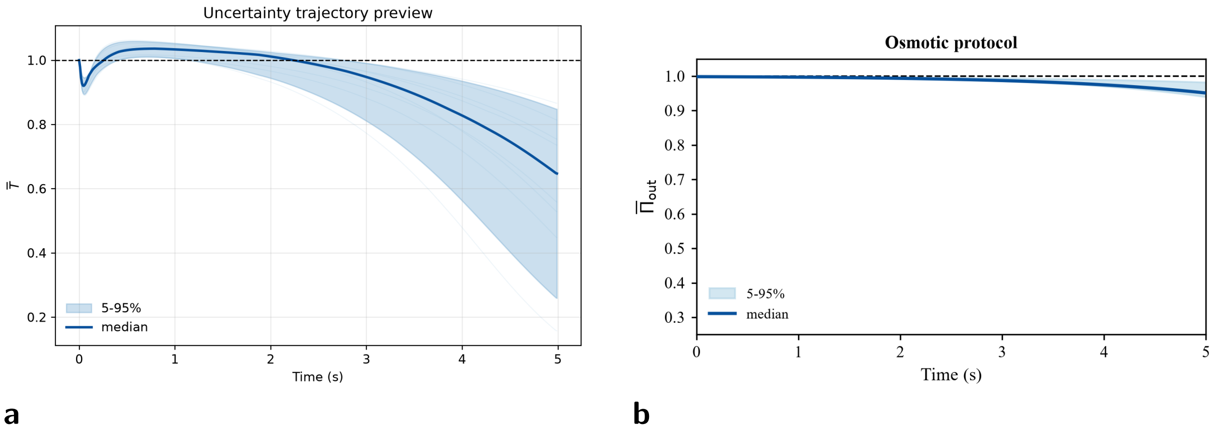


Figure S8: Conditional uncertainty analysis. (a) Uncertainty trajectory of the normalized edge traction $\overline{T}$ during the hypotonic shock window. The solid line denotes the median prediction and the shaded band the 5th–95th percentile range across sampled dynamic parameter sets. (b) Osmotic protocol used in the uncertainty analysis.

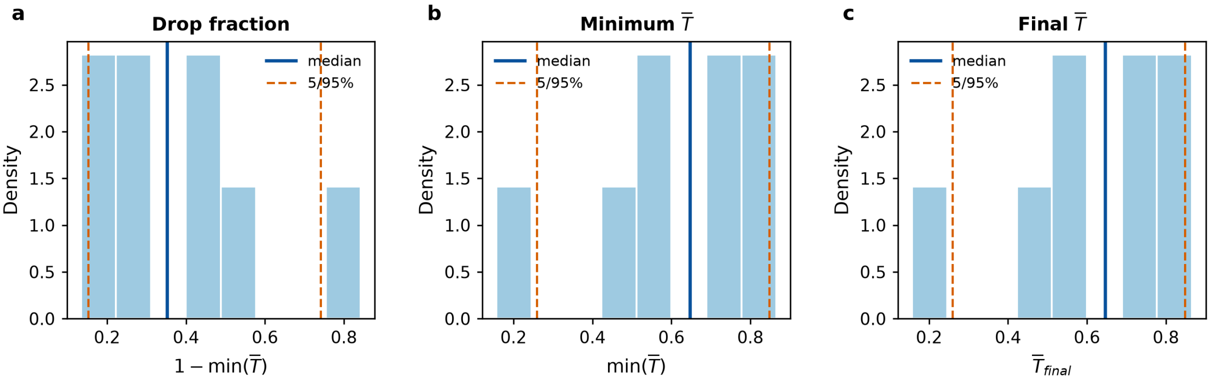


Figure S9: Uncertainty distributions of hypotonic traction-loss metrics. (a) Drop fraction $1 - \min(\overline{T})$. (b) Minimum normalized edge traction $\min(\overline{T})$. (c) Final normalized edge traction $\overline{T}_{\text{final}}$. The solid and dashed lines mark the median and 5th/95th percentiles, respectively.

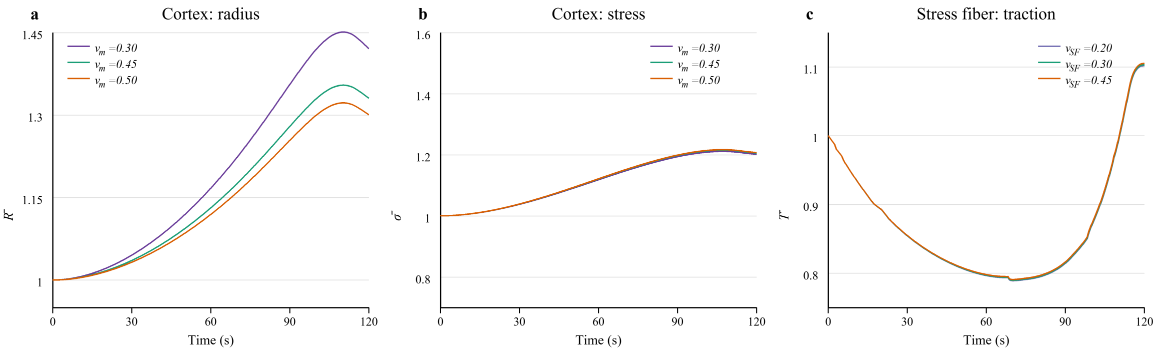


Figure S10: Poisson-ratio control related to volume-dependent mechanical response. This control separates the effects of the cortical Poisson ratio and the stress-fiber Poisson ratio. Panels (a,b) vary the cortical Poisson ratio $\nu_m$ in the cortical stress calculation and report the normalized radius $\overline{R}$ and normalized cortical stress $\overline{\sigma}$. Panel (c) varies the stress-fiber Poisson ratio $\nu_{\text{SF}}$ in the traction calculation and reports the normalized edge traction $\overline{T}$.

#### S4 Model Parameters

Table S2: Model parameters, values, and source notes used in the model calculations.

| Symbol | Description | Value | Unit |
|---|---|---|---|
| $E_m$ | Cortical modulus | 1230 (stiff); 600 (soft), AFM-measured | Pa |
| $d$ | Cortical thickness | 0.5 (1) | $\mu$m |
| $\nu_m$ | Cortical Poisson ratio | 0.5 (1) | – |
| $\nu_{\mathrm{SF}}$ | Stress-fiber Poisson ratio | 0.3 (2) | – |
| $\nu_s$ | Substrate Poisson ratio | 0.5 (3) | – |
| $\sigma_{\mathrm{act}}$ | Active cortical stress | 100 (1) | Pa |
| $v_{\mathrm{p}}$ | Polymerization velocity | 500 (4–6)[a] | $\mathrm{nm\,s^{-1}}$ |
| $\varepsilon_a(0)$ | Initial active strain of stress-fiber network | $-0.05$ (7) | – |
| $\Pi_{out,0}$ | Initial external osmotic pressure | 0.5 (8) | MPa |
| $\Delta\Pi_c$ | Critical osmotic pressure difference of ion pump | 30 (1) | GPa |
| $\alpha$ | Water-transport rate constant | $2.5 \times 10^{-10}$ (1)[b] | $\mathrm{m\,Pa^{-1}\,s^{-1}}$ |
| $\beta$ | MS-channel ion-flux rate constant | $5.0 \times 10^{-11}$ (9, 10)[a] | $\mathrm{mol\,m^{-2}\,s^{-1}\,Pa^{-2}}$ |
| $\gamma$ | Ion-transporter rate constant | $1.0 \times 10^{-18}$ (9, 10)[a] | $\mathrm{mol\,m^{-2}\,s^{-1}\,Pa^{-1}}$ |
| $\sigma_c$ | Channel gating stress threshold | 1000 (1)[a] | Pa |
| $\sigma_s$ | Channel saturation stress threshold | 3000 (1)[a] | Pa |
| $E_s$ | Substrate Young's modulus | 1000 (11)[a] | Pa |
| $R_c$ | Radius of force application by a single bond | 7 (12) | nm |
| $\rho_0$ | Initial adhesion density | $2 \times 10^{16}$ (3)[a] | $\mathrm{m^{-2}}$ |
| $k^0_{\mathrm{catch}}$ | Unloaded catch-bond dissociation rate | 3 (3) | $\mathrm{s^{-1}}$ |
| $k^0_{\mathrm{slip}}$ | Unloaded slip-bond dissociation rate | 0.01 (3) | $\mathrm{s^{-1}}$ |
| $k^0_{\mathrm{on}}$ | Unloaded association rate | 0.033 (13) | $\mu\mathrm{m^2\,s^{-1}}$ |
| $x_{\mathrm{catch}}$ | Catch-bond length scale | 8 (3) | Å |
| $x_{\mathrm{slip}}$ | Slip-bond length scale | 4 (3) | Å |
| $k_b$ | Receptor-ligand bond stiffness | 0.1 (3, 13)[a] | $\mathrm{nN\,\mu m^{-1}}$ |
| $\Theta$ | Absolute temperature | 310 | K |
| $k_B$ | Boltzmann constant | $1.38 \times 10^{-23}$ | $\mathrm{J\,K^{-1}}$ |
| $R_k$ | Gas constant | 8.314 | $\mathrm{J\,mol^{-1}\,K^{-1}}$ |

*Notes.* The baseline parameter set was first constrained using AFM-informed mechanical ranges and the experimentally measured normalized cell-area recovery.

[a] The parameters were selected within literature-supported or physiologically plausible ranges during this area-recovery constraint step. Afterward, the same parameter set was kept fixed for the traction-force calculations.

[b] $\alpha$ denotes an effective hydraulic permeability coefficient for adherent cells, which can differ from the intrinsic membrane water permeability because cell–substrate confinement may hinder pericellular water exchange; we therefore set $\alpha = 2.5 \times 10^{-10}\,\mathrm{m\,Pa^{-1}\,s^{-1}}$.
